# Exposure to bullying engages social distress circuits in the adolescent and adult brain

**DOI:** 10.1101/2025.02.12.637599

**Authors:** Birgitta Paranko, Claire Garandeau, Kerttu Seppälä, Vesa Putkinen, Severi Santavirta, Jussi Hirvonen, Christina Salmivalli, Lauri Nummenmaa

## Abstract

Despite advances in understanding the psychological and social consequences of peer victimization, the immediate effects of bullying on the central nervous system remain elusive. Here we mapped the neural responses to simulated bullying in adolescents and adults and tested whether these responses are associated with real-life victimization experiences. Fifty-one adolescents aged 11–14 years, and 47 adults underwent a functional MRI (fMRI) while watching first-person videos of bullying (victimization) in the school environment, as well as neutral and positive social interactions in a similar setting. Exposure to bullying versus positive social interaction engaged the socio-emotional and threat response systems, as well as regions related to social cognition, sensory and interoceptive processing, and motor control. These responses were consistent across adolescents and adults and dependent on the current and past victimization experiences of the participants. This large-scale activation of neural systems subserving socioemotional, somatosensory, and interoceptive processing highlights how peer victimization evokes a severe stress and alarm state in the central nervous system.

## 1. Introduction

School bullying, defined as repeated aggression against a peer in a more vulnerable position, is prevalent worldwide, with almost one-third of students reporting being bullied by their peers at least once in the past month (1). Victimization by bullying is associated with increased psychological symptom load in a dose-dependent manner (2). Peer victimization is a precursor of psychiatric problems, including anxiety and depression (3), somatic symptoms such as headache and abdominal pain (4), as well as suicidal thoughts (5). In adults, large-scale fronto-temporal and limbic brain networks coordinate the processing of socioemotional information to adjust behavioural priorities during both positive and negative social interactions (6, 7). These networks are also altered in children exposed to psychological stress (8). However, it remains unresolved i) how these networks acutely respond to peer victimization during adolescence and ii) whether adolescents’ response profiles differ from those of adults.

The literature on acute neural responses during bullying experiences is sparse, and previous studies have utilized primarily artificial and simplified experimental paradigms focusing on mapping neural responses to social exclusion and rejection. Coordinate-based meta-analyses of the widely used Cyberball game, a virtual ball game where participants experience social exclusion (9), and other social exclusion and rejection paradigms (10) have found consistent activation of the frontocortical emotion regulation system (lateral prefrontal cortex), social cognition and self-evaluative system (ventral anterior cingulate cortex, posterior cingulate cortex, medial prefrontal cortex), and the affective system (orbitofrontal cortex) in response to simulated social exclusion, suggesting that social exclusion elicits a distress response in the brain.

Cross-sectional studies further suggest that chronic peer victimization and rejection could lead to hypersensitivity to future peer rejection via stronger activation of the threat-response system. Specifically, real-life victimization and peer rejection have consistently been associated with increased amygdala and fusiform gyrus activation in response to social exclusion and negative interpersonal feedback (11–13). Neural responses to peer victimization could also differ between adolescents and adults, as sensitivity to social evaluation peaks in adolescence: Adolescents start spending more time with their friends and less time with their family members, and become increasingly concerned with their status among peers (14, 15). However, meta-analytic evidence for such age-dependent sensitivity is weak, at least for the simulated social exclusion tasks, where surprisingly, only ventral striatal responses to exclusion were stronger in younger versus older subjects (10). Consequently, the developmental time course of the acute distress responses to bullying remains poorly characterised.

Bullying can take numerous forms, ranging from social exclusion to physical aggression and persistent verbal harassment. Such phenomena are difficult to investigate in the laboratory, and consequently, neuroimaging studies on life-like peer victimization are scarce. It is thus clear that simplistic experimental paradigms of social rejection and exclusion do not capture the full complexity of the social interactions involved in peer victimization and their developmental time course (10). This may explain why, for example, the widely used Cyberball social exclusion experience engages primarily the brain’s default mode network rather than the large-scale limbic and paralimbic emotion circuits typically engaged during emotionally evocative experiences (16, 17). Indeed, large-scale cortical and subcortical brain networks parse multiple, simultaneously occurring social perceptual features ranging from others’ intentions and actions to the subtle affective characteristics of social interaction and social hierarchy in adults (7, 18), and the extent of these networks goes significantly beyond those engaged in the simulated peer exclusion or rejection tasks (9, 10). Finally, there is reason to doubt whether findings from simplified experiments would apply to real-world bullying, as many psychological and social phenomena do not fit into tightly controlled stimulus models (19). Accordingly, to understand the neural responses to bullying, it is necessary to go beyond tightly controlled yet simplistic and artificial paradigms, and map the brain responses in naturalistic settings that resemble the complex dynamics of adolescents’ everyday social life.

### The current study

Here, we mapped the acute functional neural responses to simulated bullying. We measured haemodynamic brain activity while participants viewed first-person video clips representing realistic peer interactions in a school environment. The videos depicted varying degrees of bullying (peer victimization) and positive social interaction (kind behaviour). Such engaging social and emotional content presented in a naturalistic fashion makes them ideal for modelling life-like victimization experiences in the laboratory (18, 19). We then modelled the neural responses to the moment-to-moment experience of bullying^1^ and positive social interaction occurring in the videos. We compared the brain responses of adolescent (11–14 years old) with those of adults to understand how adolescents and adults process the experience of victimization. Finally, we tested how real-life victimization experiences are related to brain responses to simulated bullying, controlling for internalizing symptoms for both age groups and workplace victimization experiences for adults. We found that exposure to (simulated) bullying engaged the socio-emotional and sensory processing regions consistently in adolescents and adults, and these responses were dependent on the real-life victimization experiences of the participants.

## 2. Methods

### 2.1 Participants

The participants were 53 Finnish-speaking adolescents and 49 adults with normal or corrected to normal vision, with no current psychiatric conditions or medication affecting the central nervous system. Two participants from each group had to be excluded because of technical problems resulting in incomplete fMRI data. Hence, data from 51 adolescents (29 females, age range: 11–14, mean age 12.20±1.02) and 47 adults (29 females, age range: 19–39, mean age 24.02±4.38) were included in the sample. Sexes were matched between adolescent and adult samples (χ^2^(1) = 0.08, *p* = .78), and ages were matched between the sexes in the adolescent (*d* = 0.02, Welch’s *T*(46.276) = 0.09, *p* = .93) and adult samples (*d* = 0.09, Welch’s *T*(41.46) = 0.31, *p* = .76). Adolescents were recruited through their parents using social media, flyers, and University and University Hospital social media and mailing lists. Adult participants were students and personnel from the University of Turku. Adolescent participants received four movie tickets and a 3D print of their brain, and adults received 50 euros as compensation. All participants and the guardians of under-aged participants signed a written informed consent. The ethics board of the Hospital District of Southwest Finland approved the protocol, and the study was conducted in accordance with the Declaration of Helsinki.

### 2.2. Self-report measures

Participants completed the following self-report questionnaires before the scan: Sum of the multidimensional Peer-Victimization scale (21) was used to measure peer victimization in adolescents. The scale includes 16 self-reported items and 4 sub-scales (physical victimization, verbal victimization, social manipulation, and attacks on property), and has acceptable to excellent reliability (Cronbach’s α = .74–.96) (22). Participants were asked how often another pupil had done different adverse things to them over the last school year, and responses were scored on a three-point Likert-scale (0 = not at all, 1 = once, 2 = more than once). For adults, both retrospective victimization and current workplace victimization were measured. Participants were asked to report the duration of their victimization at school and outside of school before adulthood, scored from 0 (“Never”) to 4 (“throughout my school years”), and the higher score out of these two measures was used as the retrospective victimization score. For current workplace victimization, adults reported how often they had been the target of 20 different offending acts at their workplace or at their current community (0 = “Never”, 4 = “Once a week or more often”). Adolescent internalizing symptoms were measured as the sum score of the shortened 25-item version of the Revised Child Anxiety and Depression Scale (RCADS-25; Ebesutani et al., 2012). Both subscales have acceptable reliability in school-based samples (Total Anxiety α = .86, Total Depression α = .79) (23). For adults, the sum of depression and anxiety subscales from the Depression, Anxiety and Stress Scale - 21 Items (DASS-21; Lovibond & Lovibond, 1995) was used as a measure of internalizing symptoms. DASS-21 subscales have acceptable reliability (α ≥ .74) under the bifactor structure (24).

### 2.3 Experimental design for fMRI

We used naturalistic first-person video stimuli depicting various degrees of bullying, neutral, and positive social interaction targeted at the viewer (*link to be published later*). The videos were filmed in a school with child actors. Bullying behaviour consisted of physical, verbal, and relational victimization, such as being called names by peers or being left out of a peer group. Each video lasted from 20 to 87 seconds, and the same pool of actors and locations was used in all the videos (seven bullying videos and five positive social interaction videos). The intensity of bullying (offensive behaviour) and positive social interaction (kind behaviour) in the videos was rated by a separate pool of 271 Finnish-speaking adults in an online experiment. Each participant rated the intensity of either bullying or positive social interaction for a subset of four videos using a dynamic response slider. After removing bad quality data (technical problems etc.), data from 235 participants was used. This yielded ratings of 30-39 participants for each video at each time point. Data were collected on a 10 Hz frequency. To match the ratings with the fMRI data, mean values for bullying and positive social interaction were downsampled to the temporal resolution of the EPI data (3s). These ratings were used as regressors in the analysis of the fMRI data to model the neural responses to bullying. Additionally, participants rated the emotional content of the videos to map the discrete emotional consequences of bullying and prosocial interaction (See section “Emotions elicited by the stimulus videos” in the supplementary material and **Figure S1**).

In the fMRI experiment, participants were instructed to watch the first-person videos as if they were the person experiencing the events but to refrain from reacting by moving or talking. Altogether, the video stimulus lasted for nine minutes, and videos were presented in a fixed order without breaks to enable brain synchronization analyses in adolescents and adults. The video presentation was controlled with Presentation® software (Version 23.0, Neurobehavioral Systems, Inc., Berkeley, CA, www.neurobs.com). Visual stimuli were shown from a screen viewed by the participant via a mirror fixed to the head coil. Sensimetrics S14 insert earphones (Sensimetrics Corporation, USA) were used to deliver the video sounds, and sound level was adjusted individually for each participant. To verify that the fMRI participants perceived bullying (offensive behaviour) and positive social interaction (kind behaviour) in the videos as intended, they were asked to rate the bullying and positive social interaction content of the videos after the fMRI scan. They were shown a representative 7-14 second clip of each video, after which they were asked to report the perceived amount of bullying and positive social interaction with a slider ranging from “not at all” to “very much”.

### 2.4 Neuroimaging data acquisition and preprocessing

MR imaging was conducted at Turku PET Centre. Data were acquired using GE Signa 3T PET/MR scanner. A specially designed silent sequence was used for the T2 MRI to get the participants used to the scanner noise before the T1 MRI and fMRI, and participants could choose to watch either animal videos or an animated film during the structural MRIs to further increase comfort. High-resolution structural images were obtained with a T1-weighted (T1w) BRAVO sequence (1 mm^3^ resolution, TR 7.9 ms, TE 3.4 ms, flip angle 10°, 228 mm FOV, 256 × 256 reconstruction matrix). A total of 192 functional volumes (9 min 51 s) were acquired for the experiment with a T2*-weighted echo-planar imaging sequence sensitive to the blood-oxygen-level-dependent (BOLD) signal contrast (TR 3000 ms, TE 30 ms, 90° flip angle, 256 mm FOV, 128 × 128 reconstruction matrix, 250 kHz bandwidth, 2.7 mm slice thickness, 51 axial slices acquired sequentially in an ascending order). Caregivers joined the debriefing of the experiment for most of the adolescent participants, and for two adolescents the caregiver joined them in the scanner room for the duration of the structural MRIs to ensure comfort. Structural brain abnormalities that are clinically relevant or could bias the analyses were checked by a consultant neuroradiologist and no subjects had to be excluded from the sample.

The anatomical and functional imaging data were preprocessed with fMRIPrep (v21.0.0rc2) (Esteban, Markiewicz, et al., 2018; Esteban, Blair, et al., 2018), which is based on Nipype 1.6.1 (K. Gorgolewski et al., 2011; K. J. Gorgolewski et al., 2018). The T1-weighted (T1w) reference images volumes were corrected for intensity non-uniformity using N4BiasFieldCorrection (ANTs 2.3.3) (Tustison et al., 2010) and skull-stripped using antsBrainExtraction.sh workflow and OASIS30ANT as template. Brain tissue segmentation of cerebrospinal fluid, white matter, and grey matter was performed on the brain-extracted T1w image using FAST (Zhang et al., 2001) (FSL v5.0.11). Brain surfaces were reconstructed using recon-all (FreeSurfer 6.0.1) (Dale et al., 1999), and the brain mask estimated previously was refined with a custom variation of the method to reconcile ANTs-derived and FreeSurfer-derived segmentations of the cortical gray matter of Mindboggle (Klein et al. 2017). Spatial normalization to the ICBM 152 Nonlinear Asymmetrical template version 2009c (MNI152NLin2009cAsym) (Fonov et al., 2009) and FSL’s MNI ICBM 152 non-linear 6th Generation Asymmetric Average Brain Stereotaxic Registration Model (MNI152NLin6Asym) (Evans et al., 2012) was performed through nonlinear registration with the antsRegistration (ANTs v2.3.3) (Avants et al., 2008), using brain-extracted versions of both T1w volume and template. Normalization to the adult template was done for both adults and children to allow direct comparisons between the samples.

Functional data were preprocessed as follows: First, a reference volume and its skull-stripped version were generated using a custom methodology of fMRIPrep. BOLD runs were slice-time-corrected using 3dTshift from AFNI (Cox and Hyde, 1997) and motion-corrected using MCFLIRT (Jenkinson et al., 2002) (FSL v5.0.11). The preprocessed BOLD was then co-registered to the T1w image using bbregister (FreeSurfer v6.0.1) for boundary-based registration (Greve and Fischl, 2009) with six degrees of freedom. All volumetric transformations were applied in a single step using antsApplyTransforms (ANTs) and Lanczos interpolation. Independent-component-analysis-based Automatic Removal Of Motion Artifacts (ICA-AROMA) was performed on the preprocessed BOLD time-series to denoise the data non-aggressively after removal of non-steady state volumes and spatial smoothing with 6-mm Gaussian kernel (Pruim et al., 2015). The first two and last nine functional volumes were discarded to exclude the time points before and after the stimulus.

### 2.5 Whole-brain GLM data analysis

The fMRI data were analysed using SPM12 (Wellcome Trust Center for Imaging; http://www.fil.ion.ucl.ac.uk/spm). A general linear model (GLM) was fitted to the data to identify the brain regions activated, in a parametric fashion, by bullying and positive social interaction of the videos. In the first level analysis, standardized dynamic mean ratings for bullying and positive social interaction (resampled to one repetition time (TR) and convolved with the canonical Haemodynamic Response Function (HRF)) were entered as regressors into a single design matrix. The signal threshold was set to 10% of the global signal, and the MNI152NLin6Asym mask was used to exclude signals outside the brain. For each participant, voxel-wise contrast images were generated for the main effects of bullying and positive social interaction, as well as for the bullying-minus-positive social interaction contrast.

These contrast images were subjected to group-level analysis with one-sample *t*-test to identify the brain regions where the relationship between the intensity of bullying or positive social interaction and the hemodynamic activity was consistent across participants. A similar group-level analysis was conducted for the bullying-minus-positive social interaction contrast to identify the regions that responded more strongly to bullying versus positive social interaction. The analyses were run separately for adolescents and adults. To investigate the possible differences in brain responses between adolescents and adults, a two-sample *t*-test between the groups was conducted. Clusters surviving FDR-correction (25) (*q* < .05) at cluster level after voxel-level thresholding with a *p*-value of .001 are reported for the main analyses.

### 2.6 Region of interest (ROI) data analysis

For the ROI analysis, anatomical ROIs were extracted using AAL3v1 atlas (26). A subset of ROIs involved in socioemotional processing (7, 27), including amygdala, anterior cingulate cortex (ACC), caudate, dorsomedial prefrontal cortex (dmPFC), insula, middle cingulate cortex (MCC), posterior cingulate cortex (PCC), putamen, thalamus, ventrolateral prefrontal cortex (vlPFC), and ventromedial prefrontal cortex (vmPFC) was chosen due to these regions’ role in processing of emotions and social exclusion (see details in section “Regions of interest definitions” in the supplementary materials). Bilateral ROIs were used, but unilateral models were also run for the main analyses, and the results are reported in **Figure S2**.

Subject-specific mean *β*-values for each ROI were obtained from the first-level whole-brain contrast images. On the group level, one-sample *t*-test was used to derive 95% confidence intervals for the beta values within each ROI and predictor for adolescents and adults separately. FDR correction for *p*-values was applied to correct for testing of multiple ROIs within age groups and predictors. In addition, paired and FDR-corrected *t*-tests were used to compare the mean *β*-values between bullying and positive social interaction within each ROI. Effect sizes are reported as Cohen’s *d*. Similar analyses but with unpaired *t*-tests were conducted to compare regional differences between adolescents and adults for each predictor.

### 2.7 Associations of real-life peer victimization and brain responses

The association between real-life peer victimization and brain responses to bullying and positive social interaction was studied using explorative whole-brain GLM analyses. Victimization score was added as a predictor in the GLM in the second-level analyses separately for adolescents and adults. Current self-reported peer victimization at school was used for adolescents and retrospective measure of victimization duration during school years for adults, as the latter allows for studying long-term neural correlates of childhood victimization. Given that internalizing symptoms (12, 28–30) and current workplace victimization may affect neural processing of simulated victimization experiences, internalizing symptoms (both groups) and current workplace victimization (adults) were included as covariates. Before deciding on the final model, effects of age and sex (within adolescent and adult groups) were tested in a separate analysis. These factors were not found to affect the brain responses to bullying or positive social interaction, apart from minor differences between males and females for the bullying-minus-positive social interaction contrast (**Figure S3**). Hence, age and sex were not included in the final models. Clusters surviving FDR correction (25) (*q* < .05) after voxel-level thresholding with a *p*-value of .05 are reported. The analysis was also repeated at the ROI level using linear regression in the aforementioned ROIs. The ROI analysis results were FDR-corrected within a predictor to account for the testing of multiple ROIs. In the adult sample, two participants had missing values in the DASS-21 questionnaire, and hence only 45 adult participants were used in the GLM and ROI analysis for studying the effects of retrospective victimization.

### 2.8 Intersubject correlation analysis (ISC)

To examine temporal dynamics of synchronization of brain activation across individuals during bullying and positive social interaction, dynamic intersubject correlation (ISC) analysis was performed. See section “Intersubject correlation analysis” in the supplementary material for more details and **Figure S4** for results.

## 3. Results

### 3.1 Subjective ratings of bullying and positive social interaction of the videos

Results from the online experiment (adult sample) revealed that the videos contained behaviours perceived as intense bullying (offensive behaviour) and positive social interaction (kind behaviour), which varied over time (**Figure 1B**). Presence of bullying and positive social interaction were negatively correlated (Pearson *r* = −.60, *p* < .001, VIF = 1.56), and these behaviours had unique time series (**Figure 1B**).

**Figure 1.**
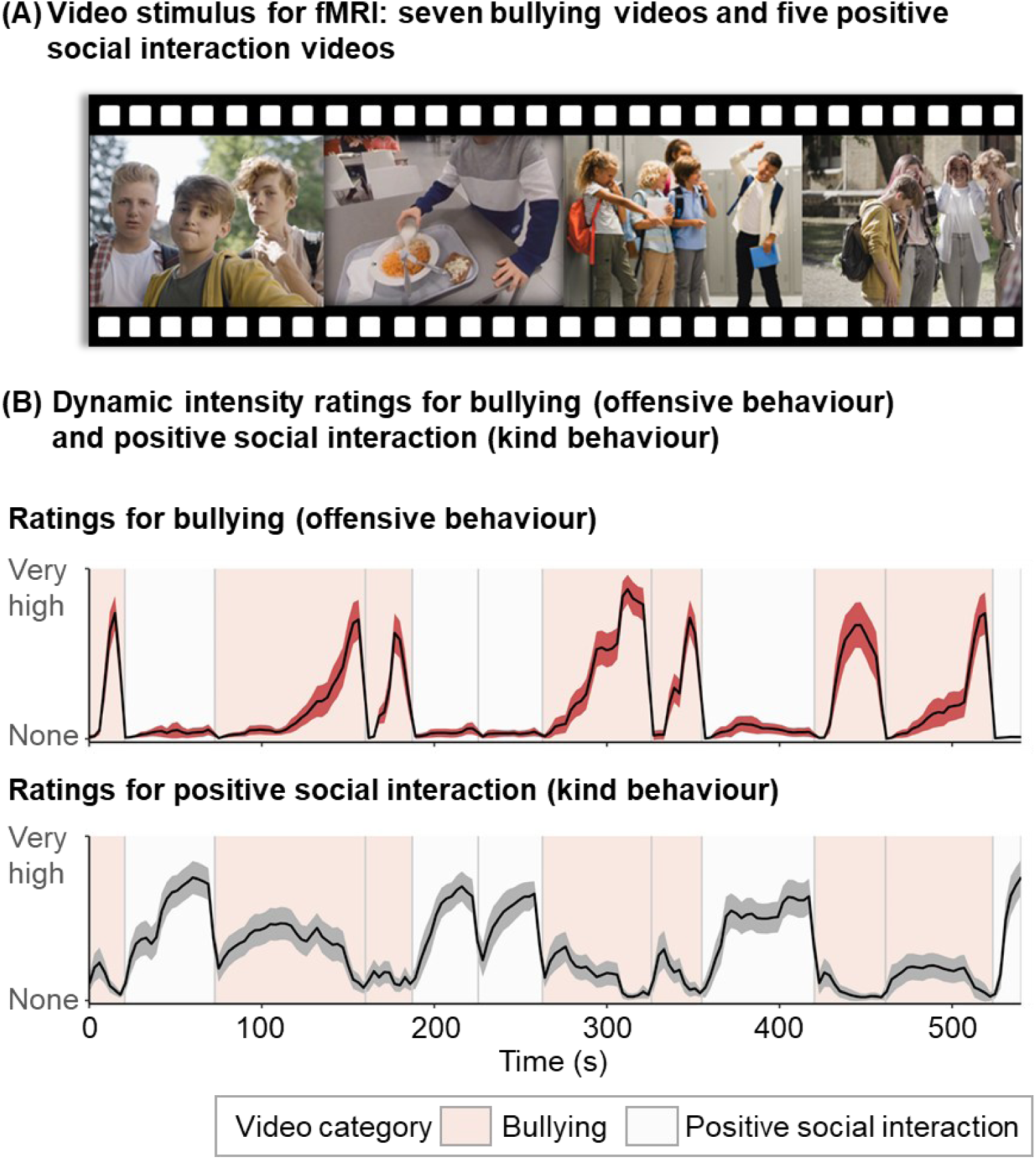
Experimental design. A) Twelve first-person videos (total duration nine minutes) were used to simulate experiences of acute bullying during fMRI. B) Mean dynamic intensity ratings with 95% confidence intervals for bullying (offensive behaviour) and positive social interaction (kind behaviour) for each video (n_raters_=235). These ratings were subsequently used as regressors in the fMRI experiment.

### 3.2 Self-reports

Self-report measures and differences between males and females are reported in **Table 1**. There were no statistically significant differences in the self-report measures between males and females in adolescents or adults. Spearman correlations between questionnaires are reported in **Figure S5**.

**Table 1.** Demographics and self-report scores.

|  | All participants<br>*M (SD)* | Females<br>*M (SD)* | Males<br>*M (SD)* | Cohen's<br>*d* | 95% *CI* for *d* | *t* | *p* |
| --- | --- | --- | --- | --- | --- | --- | --- |
| Age |  |  |  |  |  |  |  |
| Adolescents | 12.2 (1.0) | 12.2 (1.0) | 12.2 (1.0) | 0.02 | [-0.58, 0.53] | 0.09 | .93 |
| Adults | 24 (4.4) | 24.2 (4.7) | 23.8 (3.9) | 0.09 | [-0.51, 0.66] | 0.31 | .76 |
| Victimization |  |  |  |  |  |  |  |
| Adolescents | 5.8 (5.2) | 5.4 (5.5) | 6.3 (5.0) | -0.18 | [-0.79, 0.36] | -0.64 | .53 |
| Adults (retrospective) | 1.2 (0.9) | 1.2 (0.9) | 1.3 (1.0) | -0.08 | [-0.76, 0.62] | -0.25 | .80 |
| Adults (current) | 5.8 (6.3) | 5.5 (4.7) | 6.2 (8.4) | -0.11 | [-0.65, 0.67] | -0.34 | .74 |
| Internalizing symptoms |  |  |  |  |  |  |  |
| Adolescents | 13.9 (7.7) | 15.5 (8.0) | 11.7 (6.7) | 0.51 | [-0.03, 1.10] | 1.83 | .07 |
| Adults | 4.3 (3.0) | 4.4 (2.4) | 4.2 (4.0) | 0.04 | [-0.59, 0.83] | 0.11 | .91 |

*Note.* Adolescent victimization = Multidimensional Peer-Victimization scale sum score (21) (range: 0–32), adult retrospective victimization = retrospective duration of victimization before adulthood (range: 0–4), adult current victimization = workplace victimization sum score (range: 0–80), adolescent internalizing symptoms = RCADS-25 sum score (23) (range: 0–75), adult internalizing symptoms = sum of Anxiety and Depression subscales from DASS-21 (31) (range: 0–42).

### 3.3 Behavioural results for the fMRI experiment

Overall, the ratings of the fMRI participants for the videos were concordant across adolescents and adults, although adults rated the amount of bullying (offensive behaviour) slightly higher (*Mdn* = 96.43) than adolescents (*Mdn* = 91.43) in the videos categorized as bullying videos (Mann-Whitney U-test *U* =831, effect size *r* = .26, FDR-corrected *q* = .04). No other differences in ratings between groups were found. Ratings are reported in **Figure 2**.

**Figure 2.**
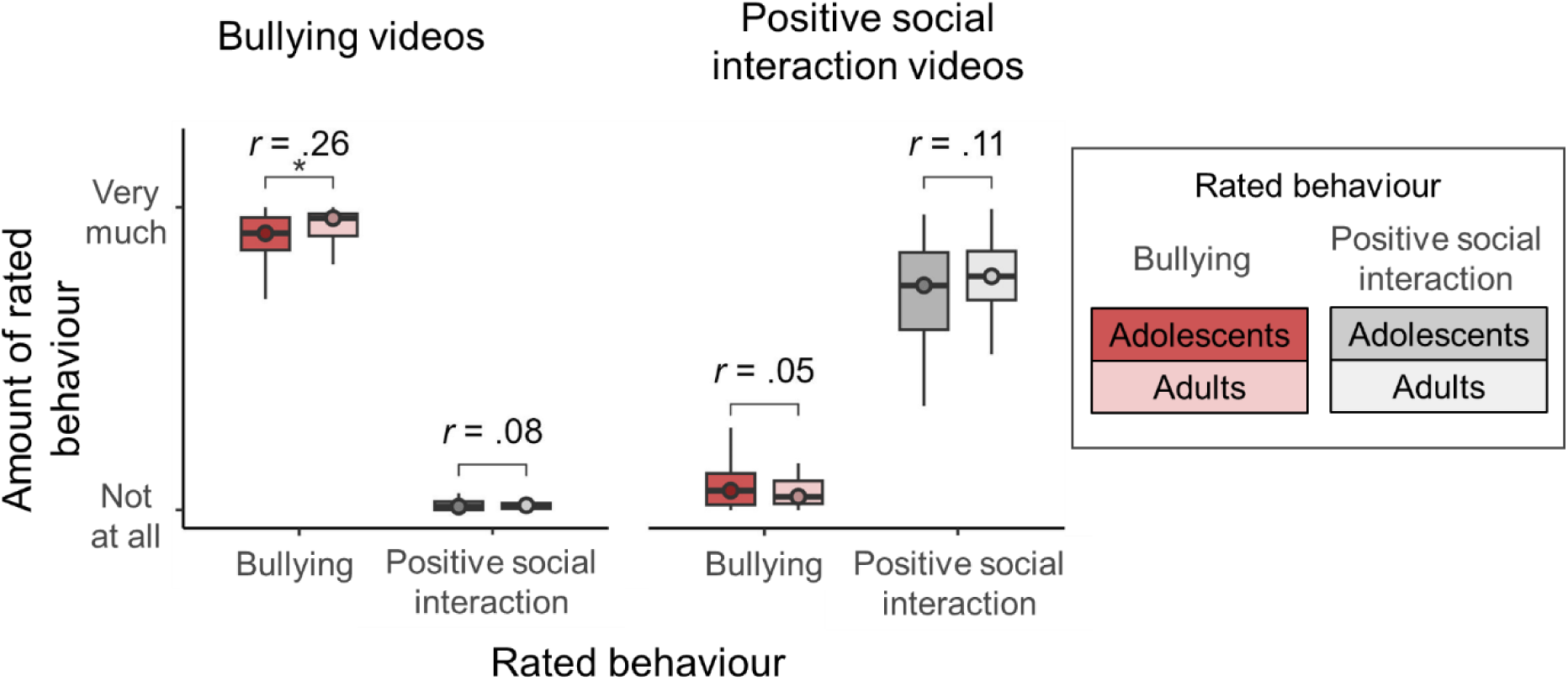
Bullying (offensive behaviour) and positive social interaction (kind behaviour) of the stimulus videos as evaluated by the fMRI participants. The horizontal line indicates the median, the lower and upper ends of the boxes indicate the lower and upper quartiles, and error bars indicate the 1.5 interquartile range. Non-parametric Mann-Whitney *U*-test was used for comparing ratings between adolescents and adults. *r* = effect size, * = FDR-corrected *q* < .05.

### 3.4 Haemodynamic responses to bullying and positive social interaction

#### 3.4.1 Whole-brain analysis

Whole-brain result maps are available on NeuroVault (https://neurovault.org/collections/BJNYACTH/.

The whole-brain GLM revealed that exposure to bullying engaged large-scale limbic and paralimbic networks in adolescents and adults (**Figure 3**). Overall, the responses were stronger and more widespread for bullying versus positive social interaction. Subcortically, bullying led to increased activity in the amygdala, caudate, hippocampus, insula, and putamen in both age groups. Cortically, the effect of bullying was particularly consistent in areas processing visual and auditory information such as lateral parts of the occipital cortex and superior temporal cortex, as well as in the fusiform gyrus. Dorsomedial and right ventrolateral parts of prefrontal cortex (dmPFC, vlPFC) were activated for both groups in response to bullying, and for adolescents, the activation extended to parts of the anterior and mid-cingulate cortex (ACC, MCC), and dorsal orbitofrontal cortex (dOFC). In contrast, a decrease in mid- and posterior cingulate cortex (MCC, PCC) activation was observed in adults in response to bullying. In adolescents, viewing bullying was also linked to activation in the precuneus and somatosensory and motor areas: primary somatosensory cortex (S1), supramarginal gyrus (SMG), supplementary motor area (SMA), and parts of primary motor cortex (M1). Similar but spatially more restricted patterns were revealed for adults, apart from S1, where activation in response to bullying was not observed.

**Figure 3.**
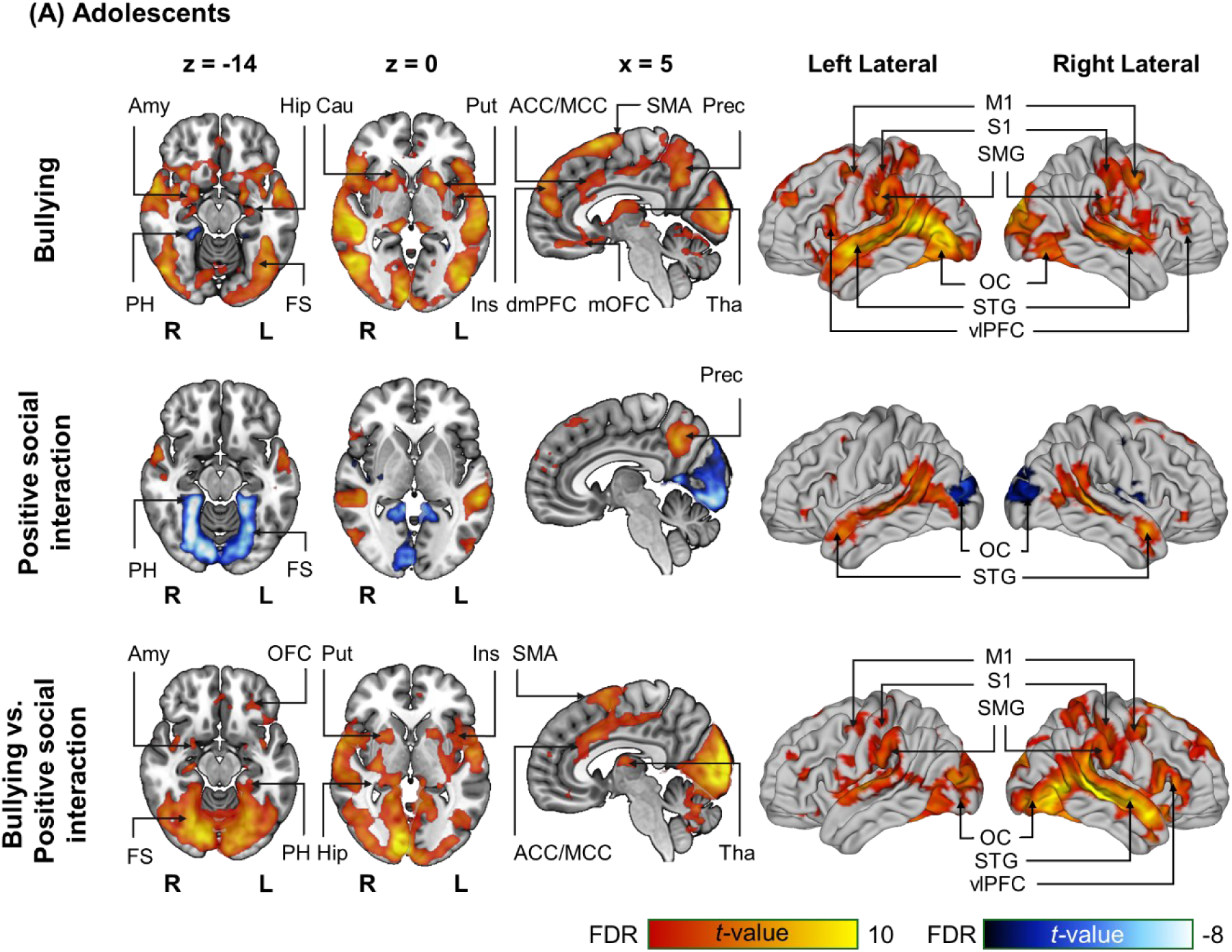
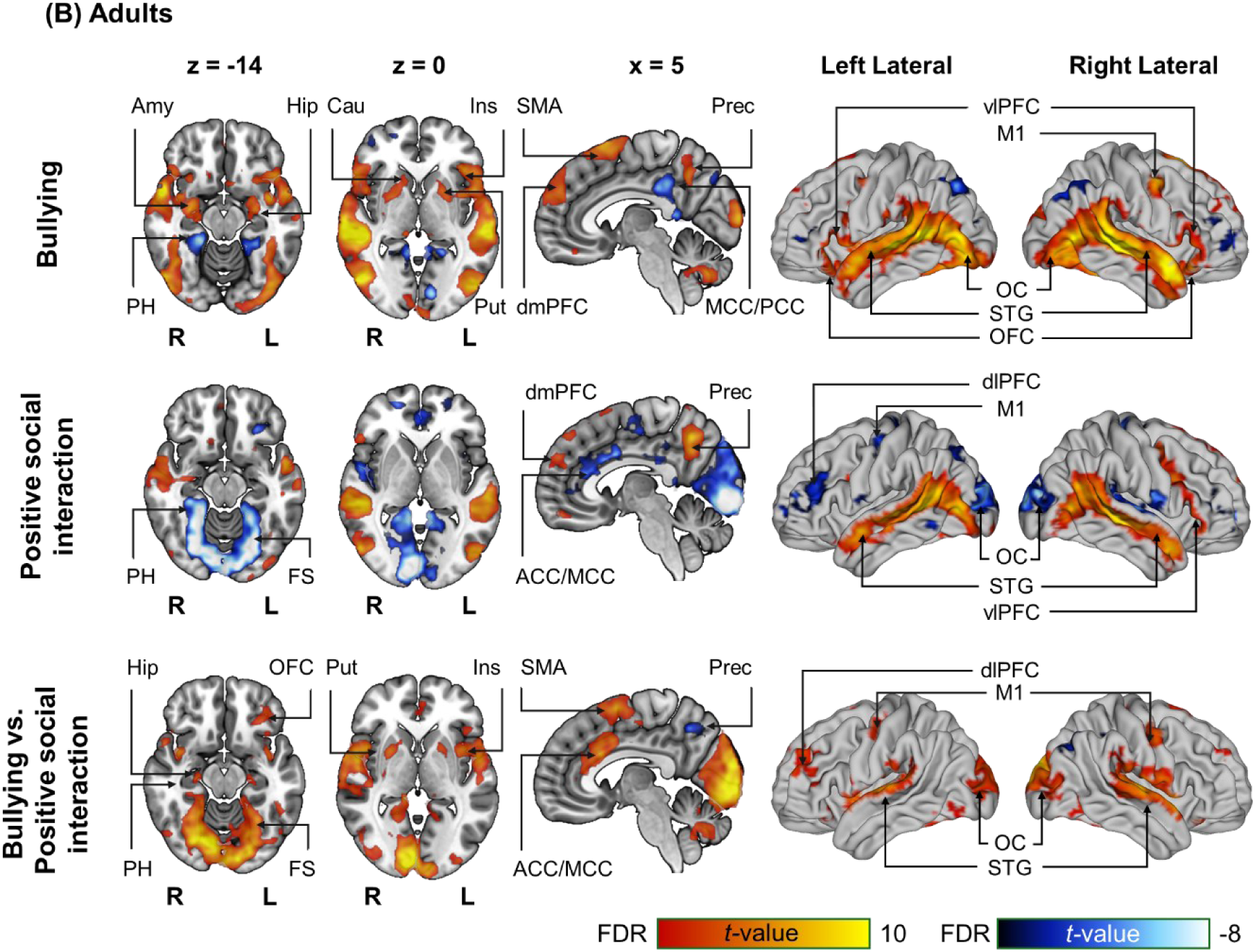
Haemodynamic responses to bullying and positive social interaction for A) adolescents and B) adults. Colour code indicates the *t*-statistic range for activations (hot colours) and deactivations (cool colours). The data are thresholded at *p* < .001 at the voxel level, and FDR corrected at *q* < .05 at the cluster level. L = left hemisphere, R = right hemisphere, ACC = anterior cingulate cortex, Amy = amygdala, Cau = caudate, dlPFC = dorsolateral prefrontal cortex, dmPFC = dorsomedial prefrontal cortex, FS = fusiform gyrus, Hip = hippocampus, Ins = Insula, M1 = primary motor cortex, MCC = mid-cingulate cortex, OC = occipital cortex, OFC = orbitofrontal cortex, PCC = posterior cingulate cortex, PH = parahippocampal gyrus, Prec = precuneus, Put = putamen, S1 = primary somatosensory cortex, SMA = supplementary motor area, SMG = supramarginal gyrus, STG = superior temporal gyrus, Tha = thalamus, vlPFC = ventrolateral prefrontal cortex.

For positive social interactions, both activation and deactivation were observed for both age groups. Activation increased in the precuneus, superior temporal gyrus, lateral occipital cortex, and restricted parts of the dmPFC and vlPFC. In contrast, deactivation was observed in the superior and medial occipital cortex, fusiform gyrus, parahippocampus, and primary motor cortex in both age groups, as well as in the left dorsolateral PFC (dlPFC) and ACC/MCC in adults.

When responses to bullying and positive social interaction were contrasted directly with each other, responses to bullying were stronger than to positive social interaction in broadly similar regions than for bullying alone, but the activation clusters were more concise. Subcortical activation was increased for bullying versus positive social interaction in the hippocampus, insula, and putamen in both groups. Amygdala activation remained significantly increased for bullying only in the adolescent group. Cortical activation was increased in the dorsal ACC and MCC, medial and superior occipital cortex, fusiform gyrus, superior temporal gyrus, SMG, SMA, M1, S1, and parts of the lateral PFC in both age groups.

#### 3.4.2. ROI analysis

The overall pattern of results in the whole-brain analysis was mostly replicated in the ROI analysis, where amygdala, insula, putamen, caudate, and vlPFC showed statistically significant responses to bullying in both age groups (FDR corrected *q* < .001, **Figure 4**). Additional responses were observed in the thalamus, ACC MCC, and dmPFC in adolescents but not in adults. No significant responses to positive social interaction were observed in adolescents, whereas in adults, deactivation was observed in response to positive social interaction in the ACC and MCC.

**Figure 4.**
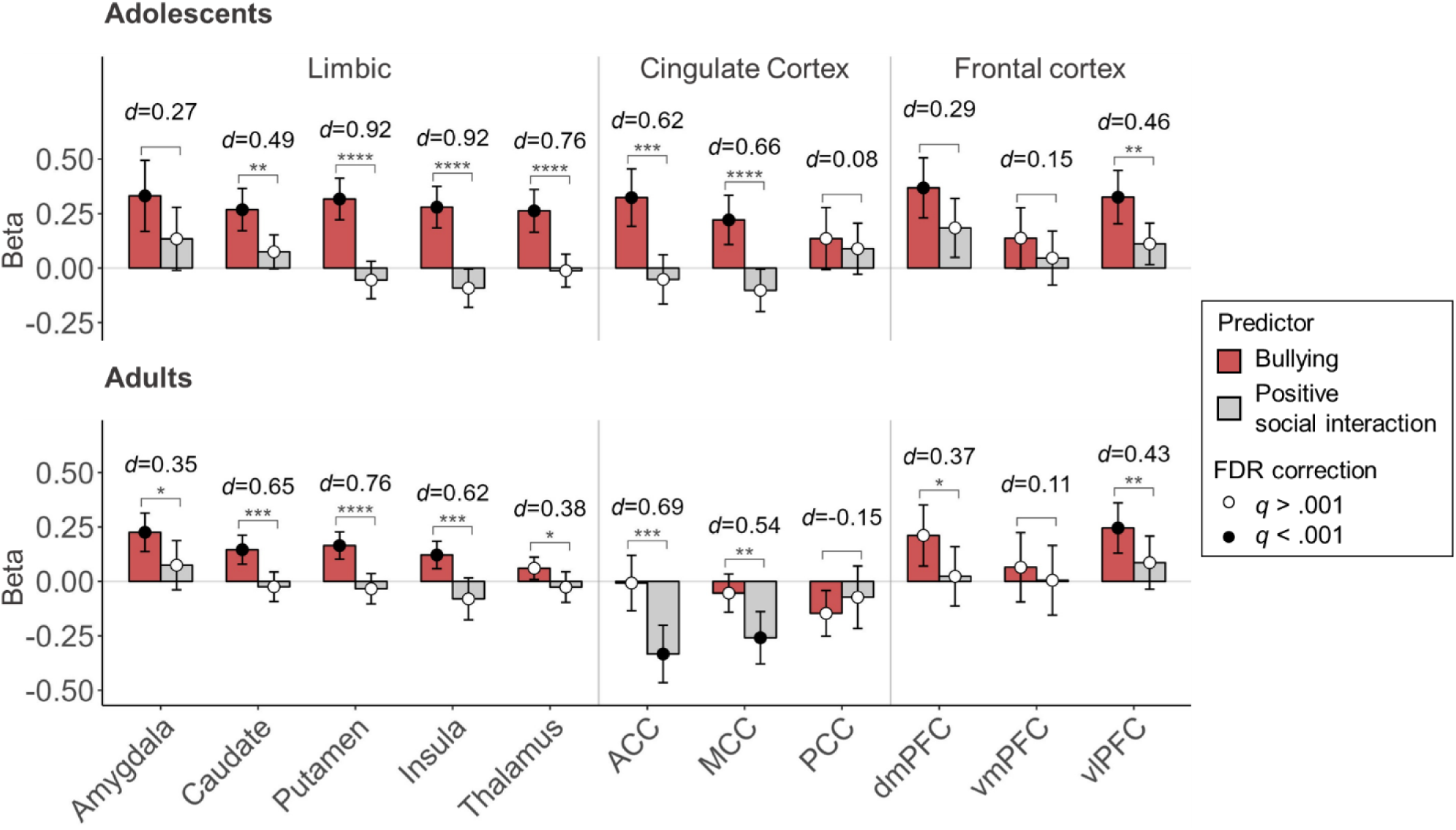
Regional effects (mean beta weights and 95% confidence intervals) for bullying and positive social interaction for adolescents and adults. Colour of the dot indicates the FDR-corrected *p*-value for one-sample *t*-test against the null hypothesis that the mean of beta-values in a specific ROI equals to zero (black: *q* < .001, white: *q* > .001, FDR corrected for multiple ROIs within age group and behaviour/condition). Asterisks denote significance levels for a paired *t*-test between bullying and positive social interaction at each ROI after FDR-correction for multiple ROIs within age group, and *d*-value indicates effect size (Cohen’s *d*) for the same test. ACC = anterior cingulate cortex, dmPFC = dorsomedial prefrontal cortex, MCC = mid-cingulate cortex, PCC = posterior cingulate cortex, vlPFC = ventrolateral prefrontal cortex, vmPFC = ventromedial prefrontal cortex. * = *q* < .05, ** = *q* < .01, *** = *q* < .001, **** = *q* < .0001.

For bullying versus positive social interaction, cortical patterns were different for ROI and full-brain analysis. The ROI analysis revealed increased activation in the vlPFC for bullying versus positive social interaction for both age groups, and an increase in dmPFC activation for adults, while in the whole-brain analysis clear vlPFC activation was only seen in adolescents, and no dmPFC activation was observed. For the cingulate cortex, the results were in line with the whole brain analysis; activation in response to bullying was higher than to positive social interaction in ACC and MCC for both age groups.

### 3.5 Comparison between adolescents and adults

Most responses to viewing bullying and positive social interaction were consistent across adolescents and adults, with spatial Pearson correlations between adolescent and adult result maps exceeding *r* = .78 (*r*_Bullying_= .78, *r*_Positive_ = .84, *r*_BullyingVsPositive_ = 0.80). Viewing bullying elicited stronger brain responses in adolescents in comparison to adults in ACC/dmPFC, MCC, fusiform gyrus, M1, S1, superior parietal cortex, and SMG, as well as areas processing visual information (cuneus, calcarine sulcus, and lingual gyrus) (**Figure S6**). For positive social interaction, adolescents’ responses were stronger than adults’ in the small parts of dlPFC and PCC. The only difference between the age groups for bullying versus positive social interaction was observed in the crossing of M1/S1, in which the difference between bullying and positive social interaction was larger in adolescents in comparison to adults.

ROI-level analysis revealed stronger responses to bullying for adolescents in comparison to adults in the insula, putamen, thalamus, ACC, MCC, and PCC (FDR corrected *q* < .05, **Figure S7** and **Table S1**). In response to positive social interaction, a significant deactivation was observed in ACC in adults in comparison to adolescents. Higher activation in response to bullying versus positive social interaction was observed in adolescents in comparison to adults in the thalamus.

### 3.6 Associations of real-life peer victimization and brain responses

In the adolescent sample, both positive and negative associations between haemodynamic responses to bullying and self-reported peer victimization at school were observed (**Figure 5A**, initial threshold of *p* < .05 at voxel level, FDR corrected at *q* <.05 at cluster level). A positive association between peer victimization and brain responses to bullying was observed in the bilateral ACC, anterior MCC, vmPFC, OFC, and SMA, right dmPFC, anterior insula, and vlPFC, as well as left ventral striatum. In contrast, victimization experiences were related to decreased responses to bullying in parts of the bilateral posterior MCC, left dorsal striatum, insula, and STG, in addition to right precuneus and angular gyrus. In the positive social interaction contrast, activation in parts of bilateral superior parietal cortex and precuneus, as well as left S1, had a negative association with peer victimization (**Figure S8A**). Victimization was associated with stronger responses to bullying in contrast to positive social interaction in the right anterior insula and vlPFC, as well as bilateral SMA and anterior MCC (**Figure S8A**).

**Figure 5.**
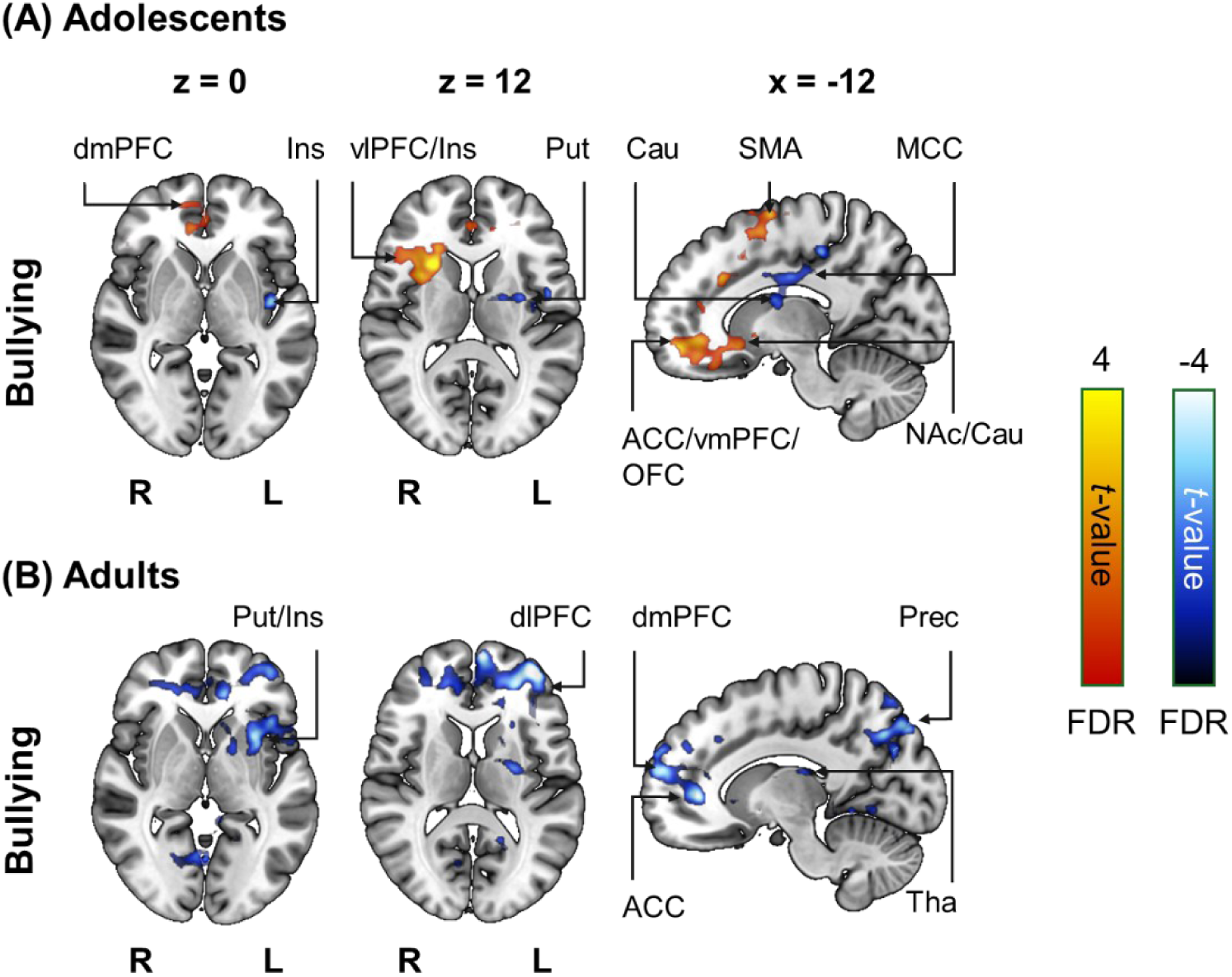
Associations between A) peer victimization in adolescents and B) retrospective peer victimization in adults and brain responses to bullying. The activation maps show *t*-values for one-sample *t*-test for the victimization scores for adolescents and adults separately, thresholded at *p* < .05 at voxel level, and FDR corrected at *q* < .05 at cluster level. Hot colours indicate positive relationship between victimization and haemodynamic responses, and cool colours indicate negative associations. Model covariates included internalizing symptoms for both groups and workplace victimization for adults. L = left hemisphere, R = right hemisphere, ACC = anterior cingulate cortex, Cau = caudate, dlPFC = dorsolateral prefrontal cortex, dmPFC = dorsomedial prefrontal cortex, Ins = insula, MCC = mid-cingulate cortex, OFC = orbitofrontal cortex, Prec = precuneus, Put = putamen, SMA = supplementary motor area, Tha = thalamus, vlPFC = ventrolateral prefrontal cortex.

In the adult sample, retrospective victimization was negatively associated with the haemodynamic responses to bullying in anterior parts of the left insula, dorsal striatum, thalamus, M1, and S1, and bilateral ACC, dmPFC, vlPFC, dlPFC, precuneus, and superior parietal cortex (**Figure 5B**, initial threshold of *p* < .05 at voxel level, FDR corrected at *q* <.05 at cluster level). Retrospective victimization was found to be associated with stronger activation in response to bullying in contrast to positive social interaction in the left posterior insula, posterior MCC, and superior temporal gyrus, as well as right thalamus (**Figure S8B**). ROI-level analysis results for the effects of victimization are reported in **table S2** and **S3**.

## 4. Discussion

Our main finding was that naturalistic simulated bullying activates the socioemotional distress circuits, including the amygdala, dorsal striatum, insula, hippocampus, and orbitofrontal cortex, as well as interoceptive and somatosensory cortices. These responses were observed in both adolescent and adult participants. This constellation of affective, somatosensory, and interoceptive activation patterns highlight the emotional burden caused by victimization and underlines how victimization induces a severe alarm or stress state in the central nervous system.

### 4.1 Brain responses to bullying

We observed large-scale limbic and paralimbic, as well as somatosensory and interoceptive activation when the participants were exposed to simulated bullying versus positive social interaction. This activation pattern indicates the recruitment of the fear and affective circuits during victimization (27). Activation was also observed in the temporoparietal and frontal regions that extract diverse and complex information from social interactions (7). Altogether, these data indicate a comprehensive shift in the social and affective processing in the brain during the threatening and adverse social interactions involving bullying. This is in stark contrast with the relatively focused activation patterns in response to social exclusion and rejection in simplified and artificial laboratory tasks, where effects are mostly observed around the cortical midline regions and in the default mode network (9, 10). All in all, our data underlines that being victimized evokes a severe state of alarm and stress in the brain.

Activation in the precuneus increased in response to bullying and positive social interaction in both age groups. Precuneus has been suggested to encode others’ intentions, and its activation likely reflects the encoding of the intended actions of the perpetrators (10). Activation of this region was stronger in response to positive social interaction versus bullying in adults, suggesting that experiencing victimization may acutely decrease the mental effort used for understanding the minds of others, possibly due to guiding cognitive capacities towards survival functions and emotion regulation. Meta-analyses on social exclusion have highlighted the role of ventrolateral PFC in emotion regulation. We observed increased activation of the vlPFC in response to bullying versus positive social interaction for adolescents in the whole-brain analysis, and ROI-analysis indicated a significantly stronger activation for bullying versus positive social interaction in vlPFC for both age groups. Based on these results, vlPFC recruitment during victimization was supported at least for adolescents, but overall the magnitude of the effect was small.

Viewing bullying versus positive social interaction also activated regions involved in social perception, mainly the fusiform gyrus, superior temporal cortex, and occipital cortex (7). Affective valence is one of the primary evaluative dimensions for social scenes and its processing engages large-scale temporoparietal and limbic regions (7, 18). Apart from the occipital pole, these regions have not been consistently observed in meta-analytic studies on simulated social exclusion (9, 10). The most likely reason for these discrepancies is the lack of detailed social information in the Cyberball paradigm, but also the differences in the type of social scenarios: In addition to peer rejection, our study also involved stimulus episodes pertaining to physical aggression and humiliation, which are more broadly representative of real-life bullying scenarios. Finally, motor cortical (M1 and SMA) activity increased in both groups in response to bullying versus positive social interaction, suggesting that simulated victimization induces initiation of motor actions possibly related to escape or initiation of counter-aggression in the threatening situation. Anterior insula activity together with posterior sensorimotor regions of the insula in both age groups, and cortical somatosensory regions (S1 and SMG) in adolescents, indicate that the affective experiences related to victimization also have a strong corporal and visceral component.

### 4.2 Consistent responses to bullying in adolescents and adults

In general, the responses to bullying were consistent across adolescents and adults, yet the activation patterns were more widespread in adolescents. The only differences that remained between the groups when contrasting bullying with positive social interaction were observed in the thalamus (ROI analysis) and the crossing of M1 and S1 (whole-brain analysis). These results indicate that simulated victimization may have been a more bodily or visceral experience for adolescents than for adults. The absence of more widespread differences between the age groups indicates that the acute effects of victimization remain consistent from adolescence to adulthood. A prior meta-analysis on the role of ACC in social rejection found ACC to be more activated in adults than in children in response to social rejection versus inclusion (32). However, we did not observe differences in the cingulate activation between adolescents and adults when comparing bullying with positive social interaction. Yet, the ROI analysis revealed that this lack of difference resulted from the fact that in adolescents the ACC response increased for bullying while remaining at zero for positive social interaction, whereas in adults the ACC response decreased for positive social interaction while remaining at zero for bullying, and the interaction contrast for these effects thus yields net zero effect. All in all, our results thus suggest the involvement of ACC during bullying in adolescents, whereas this region becomes disengaged in adults during potential social conflict, potentially reflecting the higher affective salience of social threats during adolescence.

### 4.3 Real-life peer victimization experiences predict neural responses to bullying

Finally, our results indicate that real-life victimization experiences during school years are associated with alterations in the affective stress and emotion regulation systems (ACC, medial and lateral PFC, insula, striatum). For adolescents, real-life victimization experiences were associated with increased activity of this emotion system in response to bullying, whereas in adults an opposite pattern was observed for retrospective victimization. In both groups, victimization-related increase in activation was observed for bullying in comparison to positive social interaction in the emotion and social cognition processing areas (anterior insula, vlPFC, and anterior MCC for adolescents, posterior insula, thalamus, STG and SMG for adults). These findings imply that real-life victimization may sensitize the circuits subserving emotions and their regulation in adolescence, potentially supporting immediate protection in the presence of social threats. This is in accordance with prior studies done in the context of social rejection (11–13). However, our results in the adult sample suggest that towards adulthood these continuous victimization experiences may in contrast desensitize the affective alarm system. Future longitudinal studies should confirm how these short and long-term activation patterns fluctuate throughout the development.

## 5. Limitations

Our stimuli depicted events from the school environment and were tailored for adolescent rather than adult participants. Despite this, adults and adolescents judged the bullying and positive social interaction in the videos similarly and the overall pattern of brain responses to bullying was consistent across age groups, indicating that the videos were sufficient in evoking an experience of victimization in adults. To allow comparisons across groups, all functional data were normalized into common adult stereotactic space. Despite anatomical differences between developmental groups, registration of adolescent brains in the common adult space is unlikely a source of bias on the functional level (33, 34). Finally, real-life victimization experiences were measured using self-report questionnaires, introducing a degree of uncertainty due to possible inaccuracy in recalling the actual experiences.

## 6. Conclusions

We conclude that exposure to life-like, naturalistic bullying acutely engages the socioemotional distress system and social processing regions of the brain in adolescents and adults. These responses are accompanied by increased activity in the somatosensory and interoceptive cortices, indicative of strong visceral and corporal components in the bullying experience. Altogether, this large-scale activation of neural systems subserving socioemotional, somatosensory, and interoceptive processing highlights the adverse and threatening nature of bullying, and reveals how it evokes an acute stress and alarm state in the central nervous system. Future longitudinal imaging studies should address how different risk and protective factors affect the way the brain reacts and adapts to sustained victimization.

## Acknowledgements

We thank all the participants and their families for participating in this study, Jinglu Chen, Taru Puustjärvi, and Turku PET Centre staff for assistance in data collection, and Mika Kurkilahti from A1 media for the stimulus videos. This work was supported by the Eino Jutikkala Fund, Finnish Governmental Research Funding (VTR) for Turku University Hospital and for the Western Finland collaborative area, Finnish Brain Foundation, Olvi Foundation, and The Paulo Foundation grants to BP, ERC Advanced Grant (2019/#884434) to CS, and ERC Advanced Grant (#101141656) to LN.

## Footnotes

1 The textbook definition of bullying includes repetitive behaviour and imbalance in power dynamics, yet the current stimuli provide only singular “snapshot” of the canonical bullying experience. We nevertheless refer to them as bullying because they are used for modelling the real-life bullying scenarios within the limits of the imaging laboratory context. Moreover, majority of adolescents define bullying as negative behaviour without including criteria of repetitive nature and power imbalance (20), thus the stimuli adhere well with the subject population’s typical definition of bullying.

