## Supplementary materials for "Exposure to bullying engages social distress circuits in the adolescent and adult brain"

### SUPPLEMENTARY MATERIAL

#### Emotions elicited by the stimulus videos

In the online experiment, videos categorized as bullying videos evoked stronger negative emotions, including fear, sadness, anger, disgust, shame, anxiety and depression, as well as surprise, in comparison to videos categorized as positive social interaction (**Figure S1**). In contrast, videos categorized as positive social interaction evoked stronger feelings of joy and pride.

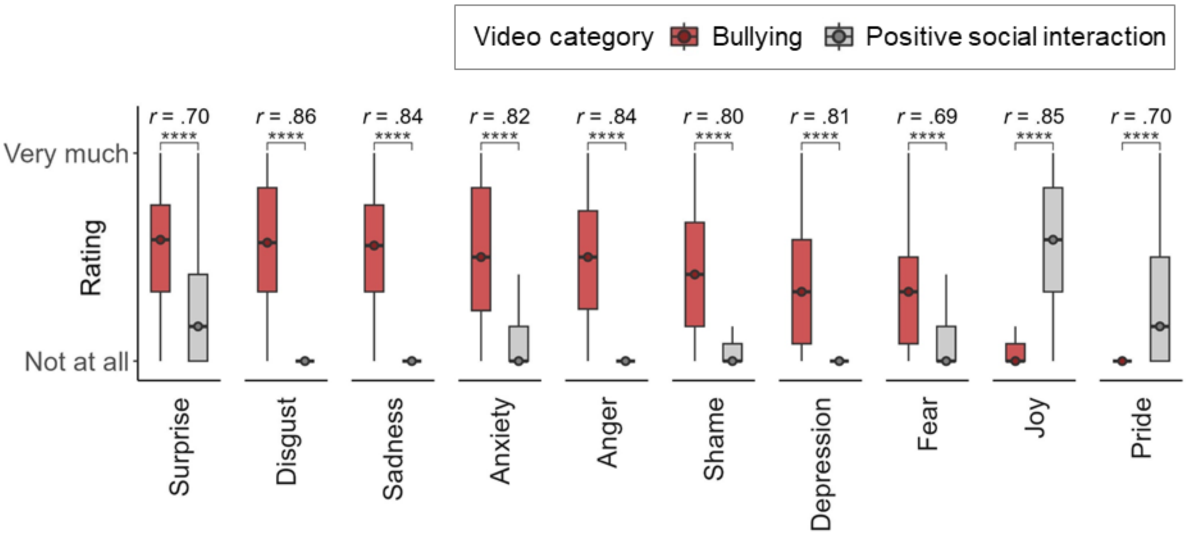


**Figure S1.** *Emotions evoked by the stimulus videos (n=212 adults from an online experiment).* Horizontal line indicates the median, lower and upper ends of the boxes indicate the lower and upper quartiles, and error bars indicate the 1.5 interquartile range. Non-parametric Wilcoxon's paired signed-rank test was used for comparing emotions evoked by the bullying and positive social interaction videos. *r* = effect size. **** = FDR-corrected *q* < .0001.

### Regions of interest definitions

Regions of interest (ROIs) were composed from the AAL3v1 atlas as follows:

| Region of interest | AAL3v1 regions |
|---|---|
| Amygdala | Amygdala |
| Anterior cingulate cortex (ACC) | ACC_pre, ACC_sub, and ACC_sup |
| Caudata | Caudate |
| Dorsomedial prefrontal cortex (dmPFC) | Frontal_Sup_Medial |
| Hippocampus | Hippocampus |
| Insula | Insula |
| Mid-cingulate cortex | Cingulate_Mid |
| Posterior cingulate cortex | Cingulate_Post |
| Thalamus | All the 12 subregions of Thalamus |
| Ventromedial prefrontal cortex (vmPFC) | Frontal_Med_Orb |
| Ventrolateral prefrontal cortex (vlPFC) | Frontal_Inf_Orb_2 |

### Unilateral ROI analysis

Confirmatory unilateral ROI analysis revealed that the observed responses to bullying and positive social interaction were bilateral to a large extent (**Figure S2**). However, in the adult sample, the effects observed in the insula and amygdala for bullying were mainly driven by the activity of the right hemisphere, whereas less consistent responses for bullying were observed on the left amygdala and insula.

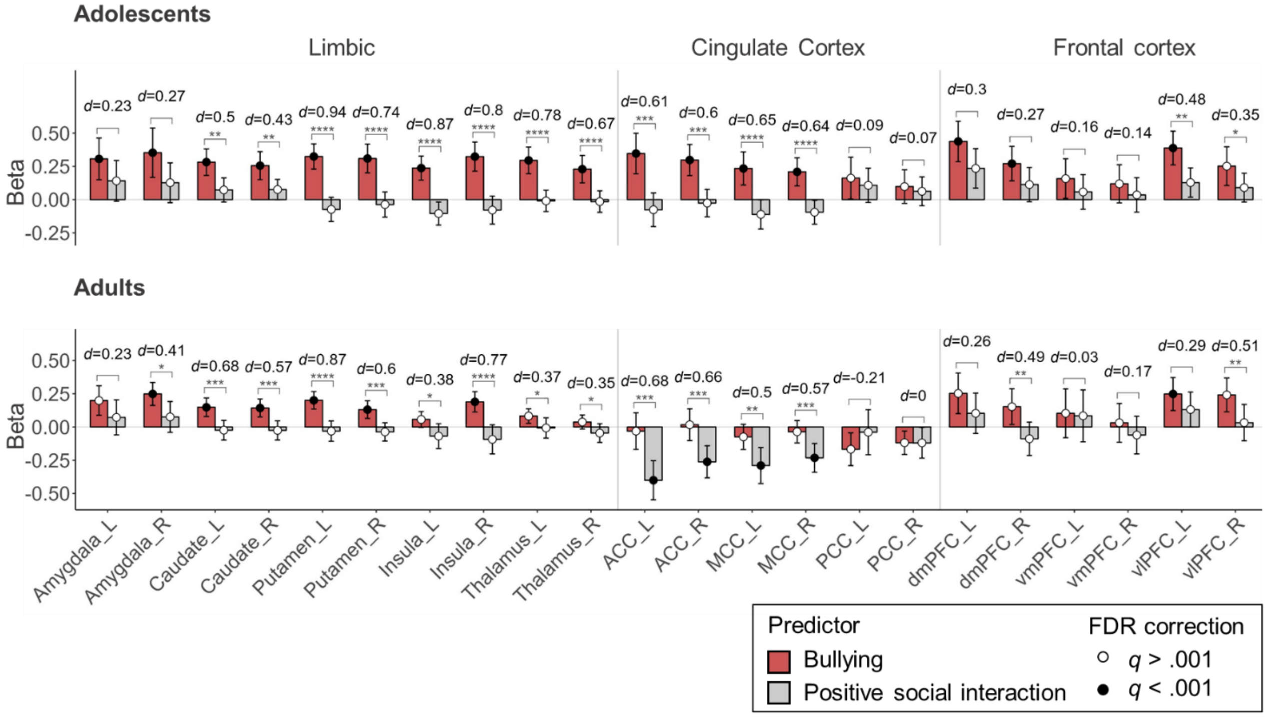


**Figure S2.** *Unilateral regional effects (mean beta weights and 95% CIs) for bullying and positive social interaction for adolescents and adults.* Colour of the dot indicates the FDR-corrected *p*-value for one-sample *t*-test against the null hypothesis that the mean of beta-values in a specific ROI equals to zero (black: $q < .001$, white: $q > .001$, FDR corrected for multiple ROIs within age group and condition). Asterisks denote significance levels for a paired *t*-test between bullying and positive social interaction at each ROI after FDR-correction for multiple ROIs within age group, and *d*-value indicates effect size (Cohen's *d*) for the same test. L = left hemisphere, R = right hemisphere, ACC = anterior cingulate cortex, dmPFC = dorsomedial prefrontal cortex, MCC = mid-cingulate cortex, PCC = posterior cingulate cortex, vlPFC = ventrolateral prefrontal cortex, vmPFC = ventromedial prefrontal cortex. * = $q < .05$, ** = $q < .01$, *** = $q < .001$, **** = $q < .0001$.

### Sex effects

Effect of sex and age was modelled for both age groups. Males were found to have higher activation for bullying versus positive social interaction in comparison to females in small regions of the SMA and precuneus (**Figure S3,** thresholded at $p < .001$ at voxel level, and FDR corrected at $q < .05$ at cluster level). Because no other effects were found for sex or age, these variables were not included in the final models.

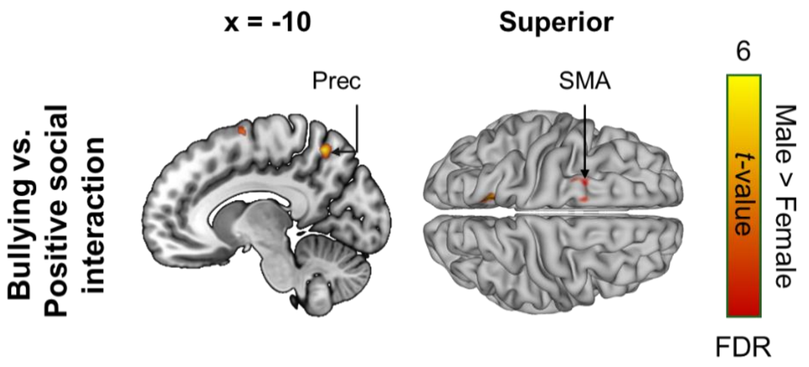


**Figure S3.** *Differences between adult males and females in haemodynamic responses to bullying versus positive social interaction.* The activation maps show *t*-values for one-sample *t*-test for sex, thresholded at $p < .001$ at voxel level, and FDR corrected at $q < .05$ at cluster level, adjusted to mean age. Positive values indicate stronger responses for males than for females. Prec = Precuneus, SMA = supplementary motor area.

### Intersubject correlation analysis

The data were detrended to remove the low-frequency scanner drift using a 240-s-Savitzky-Golay filtering (Çukur et al., 2013). Intersubject correlation analysis over the whole video stimulus was run for both adolescents and adults separately, and for their group comparison using the ISC 3.0 toolbox (Kauppi et al., 2014) with the default settings and MNI152NLin6Asym mask. Studentization of the test static was used as recommended by Kauppi et al. (2014). In addition, to map the synchronization in response to bullying, a time-window ISC with window size 10 and step size 10 was run with default settings and MNI152NLin6Asym mask, resulting in 18 thirty-second long time windows. The whole-brain correlation matrices were then analysed in SPM12 using time window matched z-standardized mean ratings for bullying and positive social interaction as regressors in GLM. Default settings for creating the GLM model were used, apart from not utilizing high pass filtering and canonical high pass filtering.

Brain activity while viewing the stimuli was most highly synchronized in the areas processing visual and auditory information, as well as the precuneus and superior parietal cortex in both adolescents and adults (FDR-corrected, $q < .001$, **Figure S4A** and **B**). The synchronization was significantly stronger in adults in comparison to adolescents in the fusiform gyrus, occipital cortex, and superior temporal gyrus (FDR-corrected, $q < .05$) (**Figure S4C**). ISC analysis indicated also stronger synchronization in adolescents in the subcortical regions (amygdala, putamen, and caudate) and cerebellum, but these results were limited to a few voxels. Time-window ISC analysis did not reveal any links between brain synchronization and the bullying content of the videos ($p < .05$, FDR corrected at cluster level at $q < .05$).

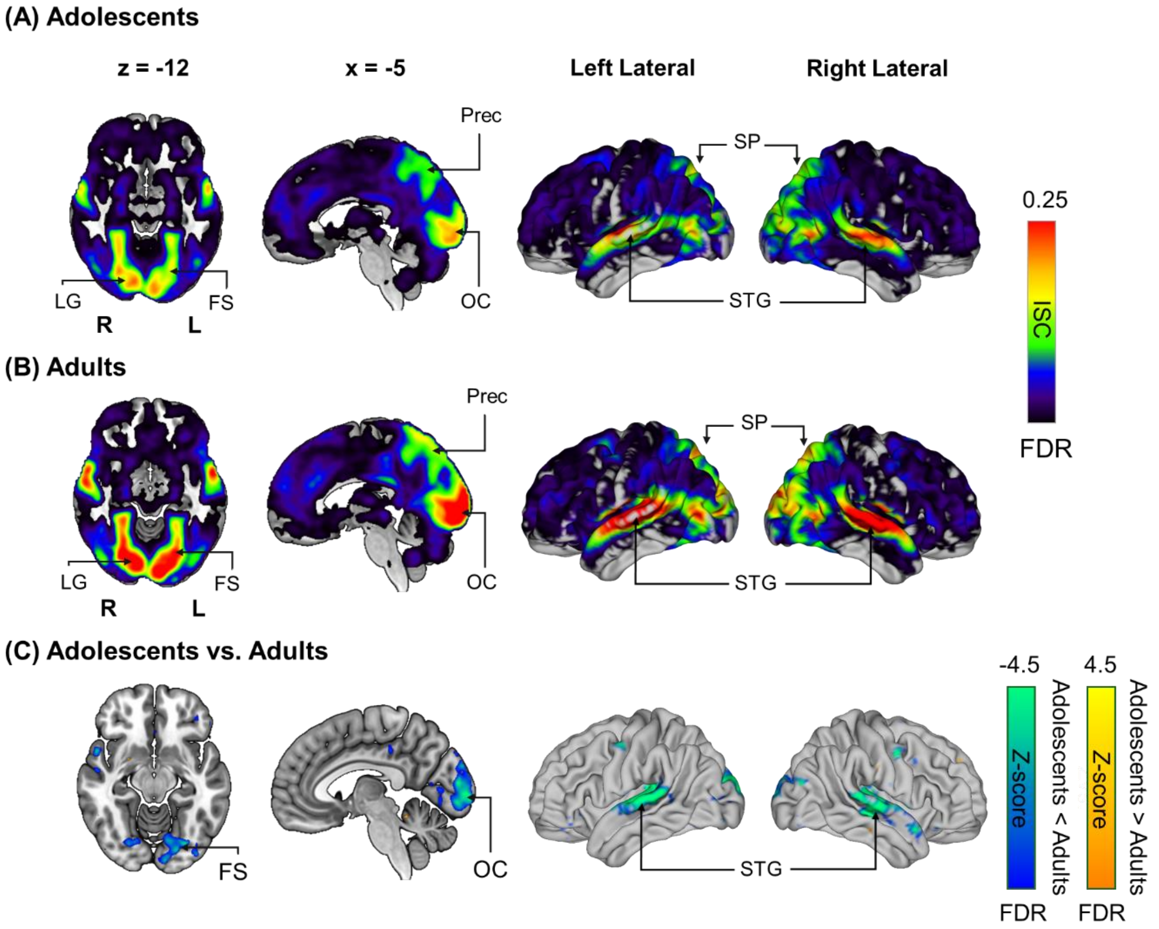


**Figure S4.** *Intersubject correlations in adults and adolescents.* Significant ISC (FDR-corrected, $q < .001$) across participants over the whole experiment for A) adolescents and B) adults. C) Significant differences in ISC between adolescents and adults over the whole experiment reported as *Z*-scores (subject-wise permutation based group-comparison, FDR-corrected, $q < .05$). L = left hemisphere, R = right hemisphere, FS = fusiform gyrus, LG = lingual gyrus, OC = occipital cortex, Prec = precuneus, SP = superior parietal cortex, STG = superior temporal gyrus.

Highest intersubject correlations were observed in the occipital and temporal regions, as well as the parietal cortex in both age groups. This accords with previous research using naturalistic movie stimulus and showing that the ISC is generally the strongest at the primary and secondary sensory regions, extending also to higher-order associative regions (Hasson et al., 2004; Santavirta et al., 2023). No association between time-window ISC and intensity of bullying was observed, suggesting that bullying behaviour does not synchronize the brain more than positive social interaction. However, considering that we observed large-scale haemodynamic activity in response to bullying in the main analysis in both age groups, the absence of ISC in the time window analysis might be explained by the short duration of the videos and the consequent low number of data points, or alternatively by intrinsic response profiles to peer victimization across viewers. In general, our data also support previous findings on age-related increases in ISC across cortical regions (Cohen et al., 2022; Moraczewski et al., 2018), pointing towards a more functionally specific organization of the cortex towards adulthood. More specifically, we observed higher ISC in adults in the fusiform, medial occipital cortex, and in the superior temporal gyrus, indicating an age-related development of regions essential for social cognition.

### Self-report correlations

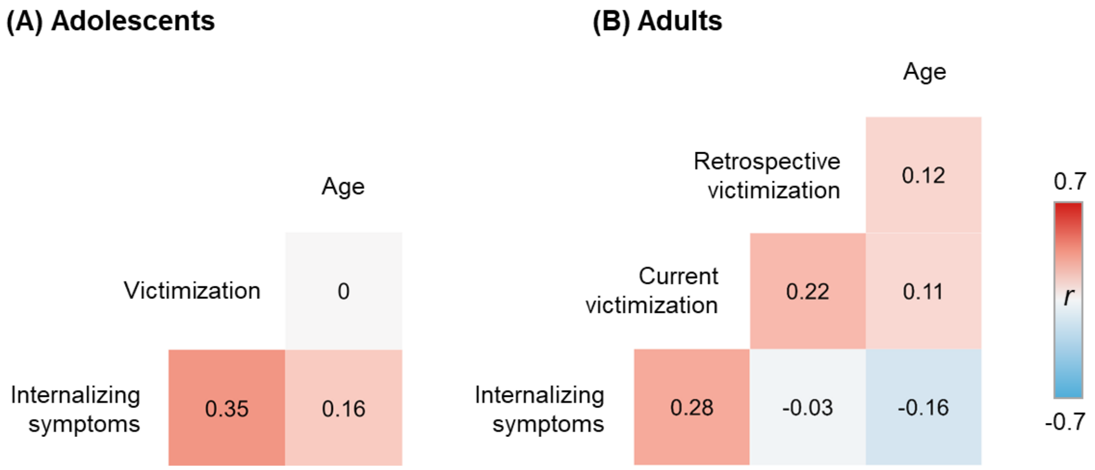


**Figure S5.** *Spearman correlations between self-report scores in A) adolescents and B) adults.* Adolescent victimization = Multidimensional Peer-Victimization scale sum score (Mynard & Joseph, 2000), adolescent internalizing symptoms = RCADS-25 sum score (Ebesutani et al., 2012), adult retrospective victimization = retrospective duration of victimization before adulthood, adult current victimization = workplace victimization sum score, adult internalizing symptoms = sum of Anxiety and Depression subscales from DASS-21 (Lovibond & Lovibond, 1995).

Comparison between adolescents and adults

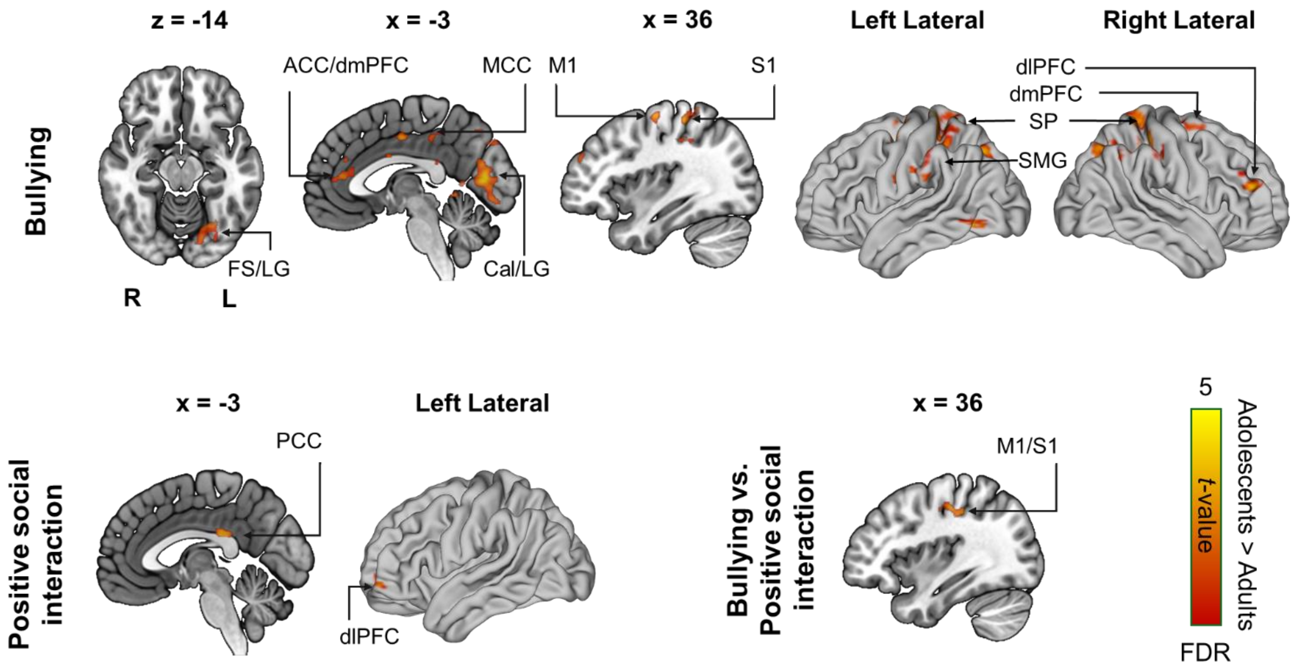


**Figure S6.** *Differences in haemodynamic responses to bullying, positive social interaction, and bullying versus positive social interaction between adolescents and adults.* The activation maps show *t*-values from two-sample *t*-test, thresholded at $p < .001$ at voxel level, and FDR corrected at $q < .05$ at cluster level. Positive values indicate stronger responses for adolescents in comparison to adults. L = left hemisphere, R = right hemisphere, ACC = anterior cingulate cortex, Cal = calcarine sulcus, Cun = cuneus, dlPFC = dorsolateral prefrontal cortex, dmPFC = dorsomedial prefrontal cortex, Fs = Fusiform gyrus, Hip = hippocampus, LG = lingual gyrus, M1 = primary motor cortex, MCC = mid-cingulate cortex, PCC = posterior cingulate cortex, S1 = primary somatosensory cortex, SMG = supramarginal gyrus, SP = superior parietal cortex.

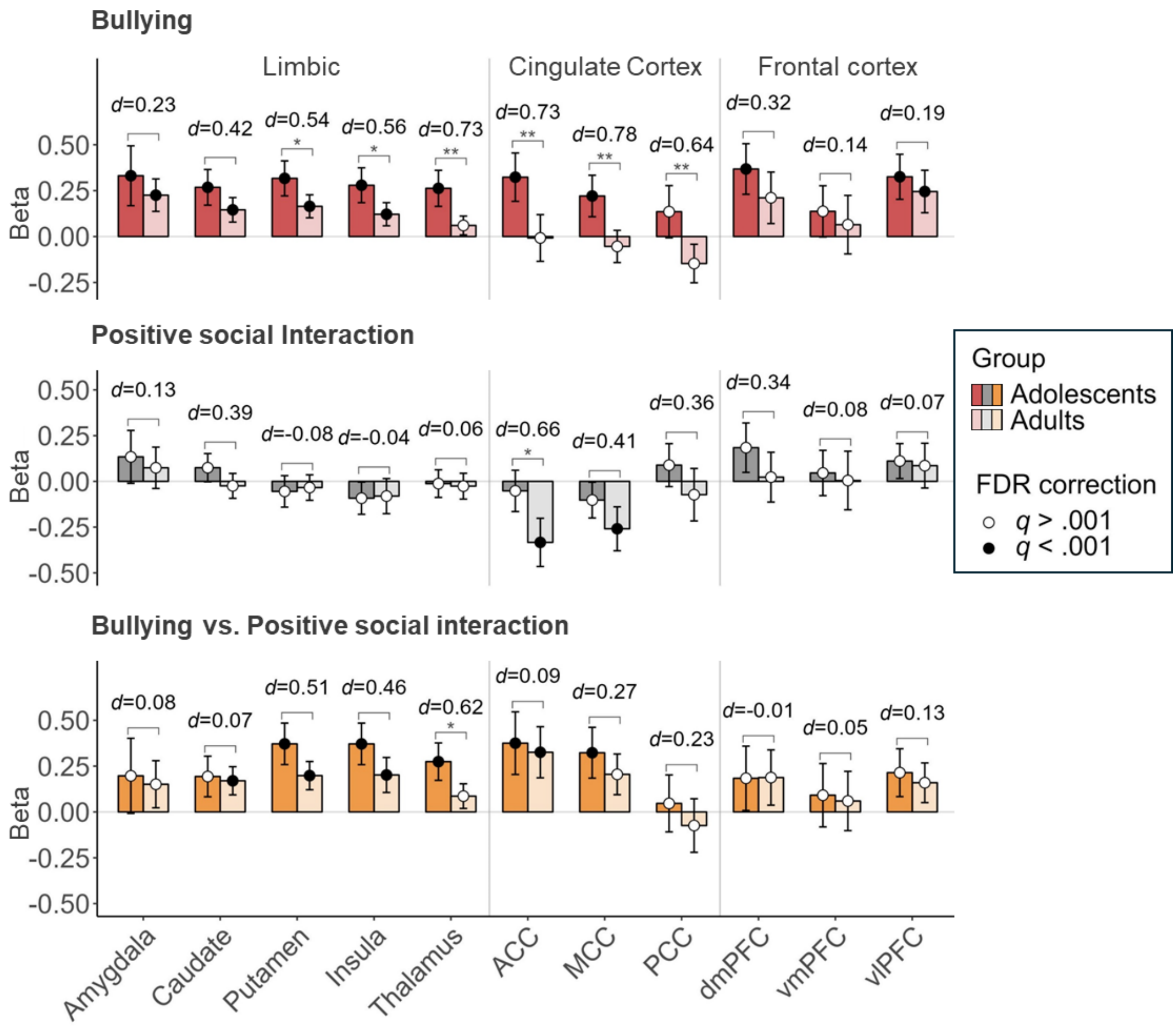


**Figure S7.** *Regional effects (mean beta weights and 95% confidence intervals) for adolescents and adults for bullying and positive social interaction, and for the bullying versus positive social interaction contrast.* Colour of the dot indicates the FDR-corrected *p*-value for one-sample *t*-test against the null hypothesis that the mean of beta-values in a specific ROI equals to zero (black: $q < .001$, white: $q > .001$, FDR correction for multiple ROI's within age group and condition). Asterisks denote significance levels for an unpaired *t*-test between adolescents and adults at each ROI after FDR-correction for multiple ROI's within condition, and *d*-value indicates effect size (Cohen's *d*) for the same test. ACC = anterior cingulate cortex, dmPFC = dorsomedial prefrontal cortex, MCC = mid-cingulate cortex, PCC = posterior cingulate cortex, vlPFC = ventrolateral prefrontal cortex, vmPFC = ventromedial prefrontal cortex. * = $q < .05$, ** = $q < .01$, *** = $q < .001$, **** = $q < .0001$.

**Table S1**

*Regional effects for adolescents and adults for bullying, positive social interaction, and bullying versus positive social interaction, and comparison between age groups.*

| ROI | Bullying | | | | | | | Positive social interaction | | | | | | | Bullying vs. Positive social interaction | | | | | | |
|---|---|---|---|---|---|---|---|---|---|---|---|---|---|---|---|---|---|---|---|---|---|
| | *M (SD)* Adolescents | *M (SD)* Adults | *t* | *df* | *d* | 95% *CI* | *q* | *M (SD)* Adolescents | *M (SD)* Adults | *t* | *df* | *d* | 95% *CI* | *q* | *M (SD)* Adolescents | *M (SD)* Adults | *t* | *df* | *d* | 95% *CI* | *q* |
| **ACC** | **0.32 (0.47)** | **-0.01 (0.43)** | **3.64** | **96.00** | **0.73** | **[0.32, 1.14]** | **.00** | **-0.05 (0.40)** | **-0.33 (0.45)** | **3.26** | **92.53** | **0.66** | **[0.25, 1.07]** | **.02** | 0.38 (0.61) | 0.33 (0.47) | 0.45 | 93.53 | 0.09 | [-0.31, 0.49] | .87 |
| Amygdala | 0.33 (0.58) | 0.23 (0.30) | 1.15 | 76.46 | 0.23 | [-0.17, 0.63] | .31 | 0.13 (0.51) | 0.07 (0.38) | 0.66 | 92.25 | 0.13 | [-0.27, 0.53] | .86 | 0.20 (0.73) | 0.15 (0.44) | 0.38 | 82.98 | 0.08 | [-0.32, 0.47] | .87 |
| Caudate | 0.27 (0.34) | 0.15 (0.23) | 2.09 | 87.19 | 0.42 | [0.02, 0.82] | .06 | 0.07 (0.27) | -0.02 (0.23) | 1.94 | 95.25 | 0.39 | [-0.01, 0.79] | .20 | 0.19 (0.39) | 0.17 (0.26) | 0.35 | 87.60 | 0.07 | [-0.33, 0.47] | .87 |
| dmPFC | 0.37 (0.49) | 0.21 (0.48) | 1.61 | 95.68 | 0.32 | [-0.08, 0.72] | .15 | 0.18 (0.48) | 0.02 (0.46) | 1.69 | 95.78 | 0.34 | [-0.06, 0.74] | .21 | 0.18 (0.62) | 0.19 (0.51) | -0.04 | 94.84 | -0.01 | [-0.40, 0.39] | .97 |
| **Insula** | **0.28 (0.34)** | **0.12 (0.21)** | **2.78** | **85.59** | **0.56** | **[0.15, 0.96]** | **.01** | -0.09 (0.31) | -0.08 (0.33) | -0.18 | 94.49 | -0.04 | [-0.43, 0.36] | .86 | 0.37 (0.40) | 0.20 (0.33) | 2.30 | 94.39 | 0.46 | [0.06, 0.86] | .09 |
| **MCC** | **0.22 (0.40)** | **-0.05 (0.30)** | **3.86** | **91.99** | **0.78** | **[0.36, 1.18]** | **.00** | -0.10 (0.35) | -0.26 (0.41) | 2.04 | 90.62 | 0.41 | [0.01, 0.81] | .20 | 0.32 (0.49) | 0.21 (0.38) | 1.34 | 93.03 | 0.27 | [-0.13, 0.67] | .51 |
| **PCC** | **0.14 (0.51)** | **-0.15 (0.36)** | **3.21** | **90.08** | **0.64** | **[0.24, 1.05]** | **.01** | 0.09 (0.42) | -0.07 (0.49) | 1.75 | 90.83 | 0.36 | [-0.04, 0.75] | .21 | 0.05 (0.55) | -0.07 (0.50) | 1.14 | 95.95 | 0.23 | [-0.17, 0.63] | .57 |
| **Putamen** | **0.32 (0.34)** | **0.16 (0.21)** | **2.68** | **85.10** | **0.54** | **[0.13, 0.94]** | **.02** | -0.05 (0.31) | -0.03 (0.24) | -0.38 | 93.42 | -0.08 | [-0.47, 0.32] | .86 | 0.37 (0.40) | 0.20 (0.26) | 2.54 | 86.64 | 0.51 | [0.11, 0.91] | .07 |
| **Thalamus** | **0.26 (0.35)** | **0.06 (0.17)** | **3.67** | **74.97** | **0.73** | **[0.32, 1.14]** | **.00** | -0.01 (0.27) | -0.03 (0.24) | 0.27 | 95.92 | 0.06 | [-0.34, 0.45] | .86 | **0.27 (0.36)** | **0.09 (0.23)** | **3.09** | **85.26** | **0.62** | **[0.21, 1.02]** | **.03** |
| vlPFC | 0.33 (0.44) | 0.25 (0.39) | 0.96 | 95.96 | 0.19 | [-0.20, 0.59] | .38 | 0.11 (0.34) | 0.09 (0.42) | 0.33 | 88.86 | 0.07 | [-0.33, 0.46] | .86 | 0.21 (0.46) | 0.16 (0.37) | 0.65 | 94.01 | 0.13 | [-0.27, 0.53] | .87 |
| vmPFC | 0.14 (0.50) | 0.06 (0.54) | 0.69 | 93.24 | 0.14 | [-0.26, 0.54] | .49 | 0.05 (0.44) | 0.00 (0.55) | 0.41 | 88.52 | 0.08 | [-0.31, 0.48] | .86 | 0.09 (0.61) | 0.06 (0.55) | 0.27 | 95.93 | 0.05 | [-0.34, 0.45] | .87 |

*Note.* Differences between age groups were tested using unpaired Welch's *t*-test and effect sizes are reported as Cohen's *d*. Statistically significant differences are boldfaced (FDR corrected $q < .05$). ACC = anterior cingulate cortex, dmPFC = dorsomedial prefrontal cortex, MCC = mid-cingulate cortex, PCC = posterior cingulate cortex, vlPFC = ventrolateral prefrontal cortex, vmPFC = ventromedial prefrontal cortex.

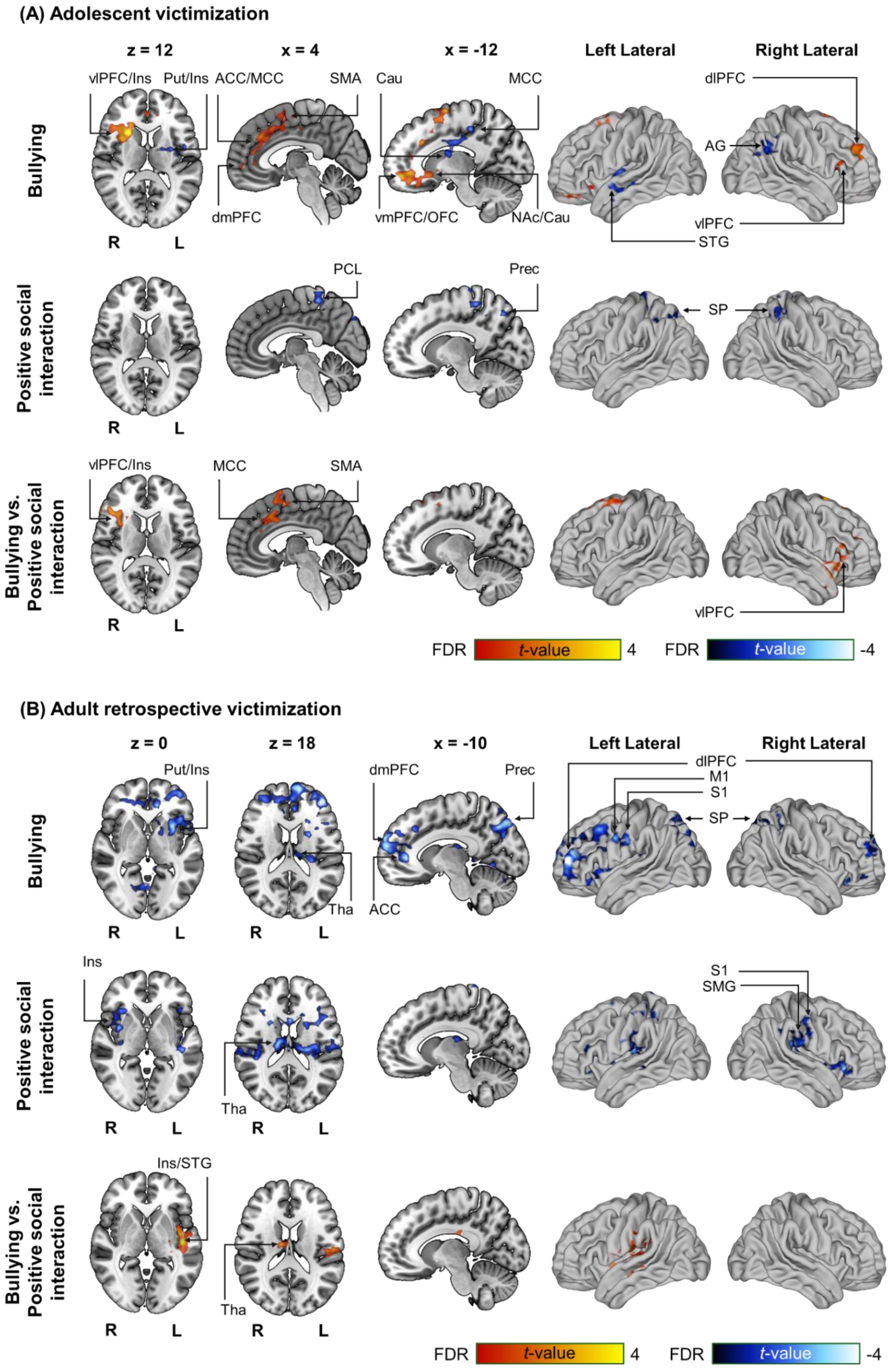


**Figure S8.** *Associations between A) peer victimization in adolescents and B) retrospective peer victimization in adults and brain responses to bullying, positive social interaction, and bullying versus positive social interaction.* The activation maps show *t*-values for one-sample *t*-test for the victimization scores for adolescents and adults separately, thresholded at $p < .05$ at voxel level, and FDR corrected at $q < .05$ at cluster level. Hot colours indicate positive relationship between victimization and haemodynamic responses, and cool colours indicate negative associations. Model covariates included internalizing symptoms for both groups and workplace victimization for adults. L = left hemisphere, R = right hemisphere, ACC = anterior cingulate cortex, AG = angular gyrus, Cau = Caudate, dlPFC = dorsolateral prefrontal cortex, dmPFC = dorsomedial prefrontal cortex, Hip = hippocampus, Ins = insula, M1 = primary motor cortex, MCC = mid-cingulate cortex, MTG = medial temporal gyrus, NAc = nucleus accumbens, OC = occipital cortex, OFC = orbitofrontal cortex, Prec = precuneus, Put = putamen, SMA = supplementary motor area, SMG = supramarginal gyrys, STG = superior temporal gyrus, S1 = primary sensory cortex, SP = superior parietal cortex, , Tha = thalamus, vlPFC = ventrolateral prefrontal cortex, vmPFC = ventromedial prefrontal cortex.

In contrast to the whole-brain analysis, ROI-level analysis did not indicate any significant effects of peer victimization on adolescents' haemodynamic responses for bullying or positive social interaction in the pre-selected regions (FDR corrected $q < .05$, **Table S2**). In adults, a negative association between retrospective victimization and brain responses to bullying was observed in the ROI analysis in the dmPFC ($R^2_{adj} = .17$, β = -0.20, 95% *CI* = [-0.34, -0.06], $t = -3.44$, $q = .01$), but not in the other preselected regions of interest nor other conditions (FDR corrected $q < .05$, **Table S3**).

**Table S2**.

*Regional effects of peer victimization experiences on adolescents' haemodynamic responses to bullying, positive social interaction, and bullying versus positive social interaction.*

| ROI | Bullying | | | | | Positive social interaction | | | | | Bullying vs. Positive social interaction | | | | |
|---|---|---|---|---|---|---|---|---|---|---|---|---|---|---|---|
| | $R^2_{adj}$ | β | 95% *CI* | t | *q* | $R^2_{adj}$ | β | 95% *CI* | t | *q* | $R^2_{adj}$ | β | 95% *CI* | t | *q* |
| ACC | .04 | 0.12 | [-0.02, 0.27] | 1.87 | .34 | -.00 | 0.05 | [-0.08, 0.18] | 0.78 | .94 | -.03 | 0.07 | [-0.13, 0.27] | 0.84 | .96 |
| Amygdala | -.01 | 0.05 | [-0.14, 0.23] | 0.72 | .96 | .01 | -0.10 | [-0.26, 0.06] | -1.51 | .94 | .05 | 0.14 | [-0.08, 0.37] | 1.74 | .65 |
| Caudate | -.01 | -0.02 | [-0.13, 0.09] | -0.36 | .96 | -.01 | -0.02 | [-0.11, 0.07] | -0.29 | .97 | -.04 | -0.00 | [-0.13, 0.12] | -0.05 | .96 |
| dmPFC | -.02 | -0.01 | [-0.16, 0.15] | -0.11 | .96 | -.01 | 0.04 | [-0.11, 0.20] | 0.67 | .94 | -.04 | -0.05 | [-0.25, 0.15] | -0.61 | .96 |
| Insula | .02 | -0.00 | [-0.11, 0.10] | -0.06 | .96 | -.02 | -0.04 | [-0.14, 0.06] | -0.66 | .94 | -.02 | 0.04 | [-0.09, 0.17] | 0.47 | .96 |
| MCC | -.04 | 0.02 | [-0.11, 0.15] | 0.34 | .96 | -.04 | -0.00 | [-0.11, 0.11] | -0.04 | .97 | -.04 | 0.03 | [-0.13, 0.18] | 0.30 | .96 |
| PCC | -.04 | 0.03 | [-0.13, 0.20] | 0.54 | .96 | -.03 | 0.05 | [-0.09, 0.18] | 0.74 | .94 | -.04 | -0.01 | [-0.19, 0.17] | -0.16 | .96 |
| Putamen | .03 | -0.02 | [-0.13, 0.08] | -0.34 | .96 | -.04 | -0.01 | [-0.11, 0.09] | -0.17 | .97 | -.01 | -0.01 | [-0.14, 0.12] | -0.13 | .96 |
| Thalamus | -.01 | 0.03 | [-0.08, 0.14] | 0.50 | .96 | -.03 | -0.01 | [-0.09, 0.08] | -0.12 | .97 | -.02 | 0.04 | [-0.08, 0.16] | 0.48 | .96 |
| vlPFC | -.03 | -0.00 | [-0.14, 0.14] | -0.05 | .96 | .00 | -0.07 | [-0.18, 0.04] | -1.08 | .94 | -.02 | 0.07 | [-0.08, 0.22] | 0.80 | .96 |
| vmPFC | .08 | 0.15 | [-0.00, 0.30] | 2.28 | .25 | -.03 | 0.02 | [-0.12, 0.16] | 0.27 | .97 | .01 | 0.13 | [-0.06, 0.32] | 1.57 | .65 |

*Note.* $N = 51$. Model statistics for victimization are adjusted to mean internalizing symptoms. All independent variables were *Z*-standardized. *CI* = confidence interval, *q* = FDR corrected *p*-value.

**Table S3**.

*Regional effects of retrospective victimization experiences on adults' haemodynamic responses to bullying, positive social interaction, and bullying versus positive social interaction.*

| ROI | Bullying | | | | | Positive social interaction | | | | | Bullying vs. Positive social interaction | | | | |
|---|---|---|---|---|---|---|---|---|---|---|---|---|---|---|---|
| | $R^2_{adj}$ | β | 95% *CI* | *t* | *q* | $R^2_{adj}$ | β | 95% *CI* | *t* | *q* | $R^2_{adj}$ | β | 95% *CI* | *t* | *q* |
| ACC | .12 | -0.12 | [-0.25, 0.01] | -2.12 | .17 | -.02 | -0.05 | [-0.20, 0.09] | -0.92 | .57 | -.01 | -0.07 | [-0.22, 0.08] | -1.08 | .77 |
| Amygdala | -.05 | 0.03 | [-0.07, 0.12] | 0.48 | .77 | -.03 | -0.06 | [-0.19, 0.06] | -1.06 | .57 | -.01 | 0.09 | [-0.05, 0.23] | 1.38 | .62 |
| Caudate | .09 | -0.04 | [-0.11, 0.03] | -0.69 | .77 | .09 | -0.06 | [-0.13, 0.01] | -1.01 | .57 | -.07 | 0.02 | [-0.07, 0.10] | 0.28 | .98 |
| **dmPFC** | **.17** | **-0.20** | **[-0.34, -0.06]** | **-3.44** | **.01** | -.04 | -0.07 | [-0.22, 0.08] | -1.15 | .57 | .03 | -0.13 | [-0.29, 0.03] | -2.05 | .45 |
| Insula | -.04 | 0.00 | [-0.07, 0.07] | 0.03 | .97 | .16 | -0.11 | [-0.20, -0.01] | -1.87 | .34 | .10 | 0.11 | [0.01, 0.21] | 1.70 | .49 |
| MCC | .04 | -0.04 | [-0.14, 0.05] | -0.76 | .77 | .02 | -0.06 | [-0.18, 0.07] | -0.98 | .57 | -.07 | 0.01 | [-0.11, 0.14] | 0.21 | .98 |
| PCC | -.06 | -0.03 | [-0.15, 0.08] | -0.57 | .77 | -.05 | -0.02 | [-0.18, 0.13] | -0.43 | .67 | -.06 | -0.01 | [-0.17, 0.15] | -0.12 | .98 |
| Putamen | .05 | -0.05 | [-0.11, 0.02] | -0.85 | .77 | .07 | -0.03 | [-0.10, 0.04] | -0.54 | .66 | -.06 | -0.02 | [-0.10, 0.07] | -0.27 | .98 |
| Thalamus | .01 | -0.00 | [-0.06, 0.05] | -0.07 | .97 | .10 | -0.03 | [-0.10, 0.04] | -0.56 | .66 | -.02 | 0.03 | [-0.05, 0.10] | 0.43 | .98 |
| vlPFC | .05 | -0.11 | [-0.24, 0.01] | -1.99 | .17 | .06 | -0.11 | [-0.24, 0.02] | -1.95 | .34 | -.07 | -0.00 | [-0.12, 0.12] | -0.03 | .98 |
| vmPFC | .03 | -0.06 | [-0.24, 0.11] | -1.10 | .74 | -.04 | -0.03 | [-0.21, 0.15] | -0.53 | .66 | -.04 | -0.03 | [-0.21, 0.15] | -0.52 | .98 |

*Note.* $N = 45$. Model statistics for retrospective victimization are adjusted to mean internalizing symptoms and workplace victimization. All independent variables were *Z*-standardized. *CI* = confidence interval, *q* = FDR corrected *p*-value. Effects with $q < 0.05$ are boldfaced.
